# Top-down and bottom-up landscape processes influence the formation of microbial partnerships in herbivorous insects

**DOI:** 10.64898/2026.05.27.728152

**Authors:** Daniel J. Leybourne, Sharon E. Zytynska, Emily A. Martin

## Abstract

Many insects form intimate associations with microbial partners, spanning a continuum from obligate mutualisms to transient associations. These microbial partners confer a range of functional benefits, including protection from natural enemies, detoxification of plant secondary metabolites, and improved fitness. The prevalence and abundance of microbial symbionts can vary between insect populations, indicating that the insect-microbe relationship is influenced by external factors. However, little is known on the factors that determine the assembly, structure, and composition of the microbiome of herbivorous insects. We posit that the factors influencing the insect-microbe relationship occur at the landscape-scale, acting through a combination of bottom-up, top-down, and indirect effects between plants, herbivores, and their natural enemies. We test this in two agro-ecosystems by sampling individual insects from three sap-feeding and one leaf-chewing species. Our results explicitly show for the first time that habitat configurational complexity and the proportion of host-plant crop in the landscape, combined with natural enemy abundance, can act as direct and indirect drivers of microbial symbiosis in herbivorous insects. This highlights the wider implications landscape habitat and ecosystem structure has on species interactions across managed agricultural landscapes.

## Introduction

Many insects form intimate associations with microbial partners, spanning a continuum from obligate mutualisms to transient associations. In herbivorous insects, microbial symbionts help their host overcome the challenges of nutrient-poor diets and protect their host from natural enemies [1]. While these microbes boost host fitness, the symbiosis often comes with a physiological or ecological cost [2, 3]. Uncovering the ecosystem-scale factors that underpin microbiome formation is crucial for designing sustainable agro-ecosystems, particularly where symbionts may disrupt beneficial services such as natural pest control.

Microbial symbionts have two major modes of transmission. Endosymbionts are predominantly vertically transmitted from parent to offspring, whereas gut-associated symbionts are largely acquired from the environment. Both types of symbionts influence insect interactions with host plants and can shape ecological networks [4]. The symbionts of herbivorous insects can contribute to metabolic flexibility; this supports the host insect in exploiting a wider range of plant resources and can also help to attenuate the production of herbivore-induced plant volatiles [5], often with cascading effects on natural enemy recruitment and foraging behaviour [6]. However, to date, little is known on the factors underlying the assembly, structure, and composition of insect microbiomes, thus reducing our ability to predict their impact at the ecosystem scale.

Aphids have become a model system for the study of insect-endosymbiont interactions [3]. Almost all aphid species form an obligatory relationship with the essential endosymbiont *Buchnera aphidicola*, and they can also host various non-essential (facultative) endosymbionts [7]. Eight common facultative species have been described, occurring as individual, co- or multi-associations [8, 9]. These endosymbionts play important ecological roles at the population and community level [3] by providing resistance against parasitoid wasp attack or fungal infection [10, 11], heat stress tolerance [12], and suppression of host plant defences [5]. Gut-associated symbionts are more important for leaf-chewing herbivores [1]. The microbiome of leaf-chewing insects is more plastic than those of sap-feeders, with transient gut microbes often transferred from the host plant to the feeding insect [13]. Despite the transient nature of the gut microbiome, it still plays a key role in detoxifying plant secondary metabolites; this is particularly important for plants with diverse biochemistries, such as the Brassicaceae [14].

Symbiont prevalence and diversity can vary within and across herbivore populations [13, 15, 16] and could be linked to the complexity and functioning of the surrounding landscape [17, 18, 19, 20, 21, 22]. Landscape configuration and the proportion of crop versus non-crop habitat are key variables for insect populations, with clear evidence that they impact resource availability for both insect herbivores and their natural enemies [21]. Landscapes with limited botanical diversity, which are dominated by monoculture crops, often suffer from increased herbivore abundance [23], while a greater proportion of seminatural habitat and greater configurational complexity support abundant natural enemy populations that suppress herbivorous insects [24]. It follows that these same variables are likely to impact insect-symbiont relationships, in turn feeding back to alter herbivore population regulation; yet, no study has comprehensively examined this.

We thus hypothesise (Fig. 1) that insect microbiomes are structured by the surrounding landscape. We posit that this occurs through bottom-up, top-down, and indirect effects between plants, herbivores, and their natural enemies, leading to the following predictions: (i) the proportion of host crops in a landscape influences herbivorous insect populations positively, through resource concentration, and positively supports microbial associations, particularly in chewing insects where the insect microbiome is influenced by diet [14]. (ii) Landscapes with more seminatural habitat support a more abundant natural enemy community, promoting herbivorous insect regulation and increasing association with protective symbionts through positive selection pressure. (iii) Crop heterogeneity provides increased botanical refugia to support greater insect populations, and the increased botanical diversity promotes insect association with detoxifying microbes. (iv) As protective phenotypes often come with fitness trade-offs [10, 3], association with protective microbes has cascading consequences that lead to seasonally-dynamic insect-symbiont associations.

**Fig. 1.**
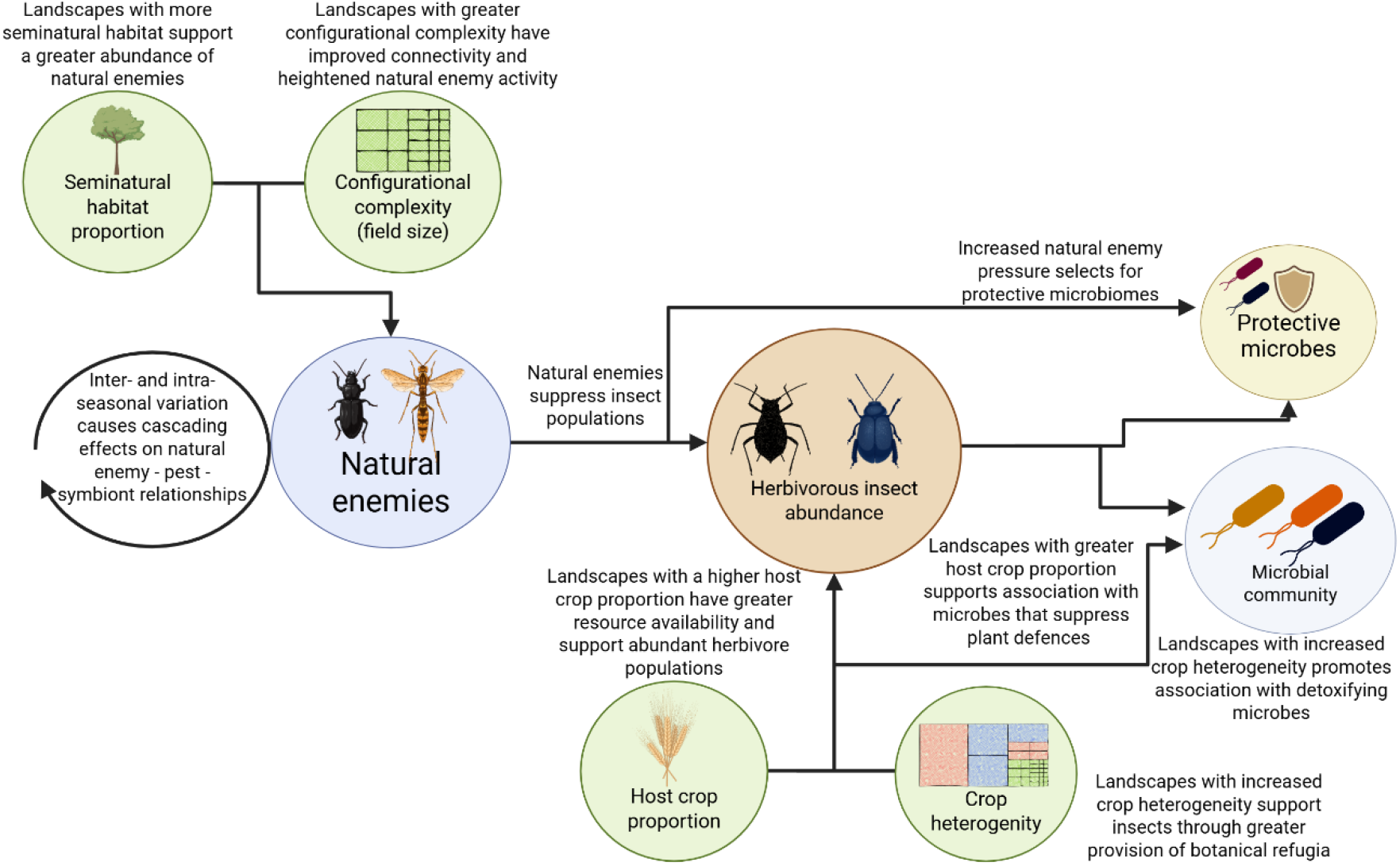
Overview of the proposed mechanisms through which the landscape might influence insect populations and the insect-microbe relationship.

Here, we used 720 field-collected aphid samples from 35 field sites and 72 cabbage stem flea beetle samples from 18 field sites to test this hypothesis. We identify several drivers of symbiont associations, indicating that top-down and bottom-up factors collectively, and interactively, influence the microbiome of herbivorous insects.

## Results

Across 792 samples taken from 53 fields (Fig. S1), we observed cryptic variation in herbivore microbiome composition and diversity both in cereal and rapeseed agroecosystems. In aphids, this included association with endosymbionts that confer resistance to natural enemies (*Regiella insecticola, Hamiltonella defensa, Fukatsuia symbiotica*, and *Ricketsiella* sp.; Fig. 2A-C). Aphid species varied in their propensity to form facultative microbial partnerships (X^2^_3_ = 11.81; p = 0.003) and in the number of endosymbionts hosted (X^2^_3_ = 41.63; p = <0.001): 60% of S*itobion avenae* and *Metopolophium dirhodum* hosted at least one endosymbiont (range 0-4 symbionts), while this dropped to just under 50% for *Rhopalosiphium padi* (range 0-5).

**Fig. 2.**
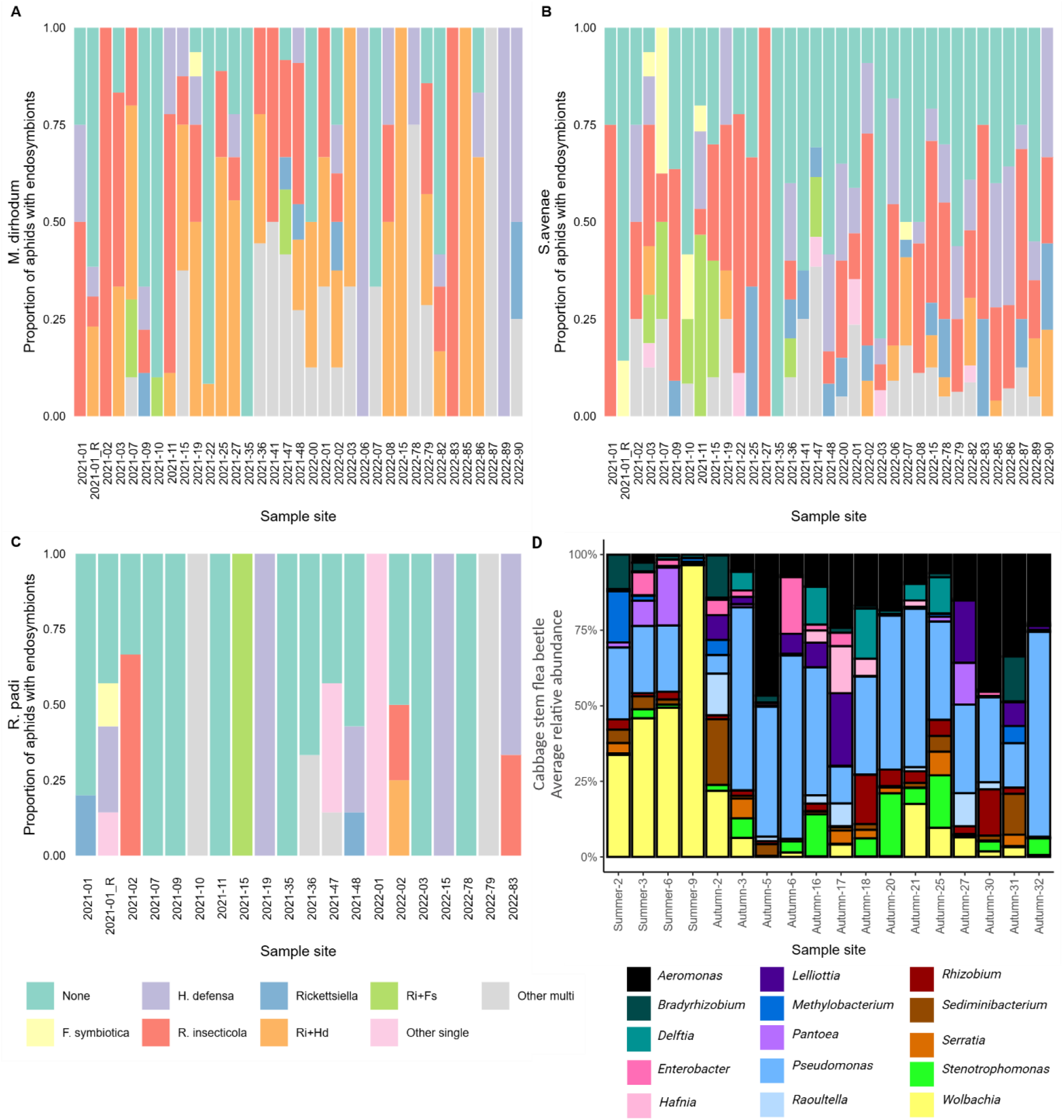
Observed endosymbiont and microbiome composition for the insect populations sampled at each site. A-C) Total proportion of individual insects hosting the respective endosymbiont species; A) *Metopolophium dirhodum;* B) *Sitobion avenae;* C) *Rhopalosiphum padi*. Endosymbionts are categorised into none (no facultative endosymbionts), *F. symbiotica, H. defensa, R. insecticola*, and *Ricketsiella spp*. in single-infections; co-infection of *R. insecticola* and *H. defensa* (Ri+Hd), co-infections of *R. insecticola* and *F. symbiotica* (Ri+Fs), single infection with other facultative species (Other single), and other co- or multi-infections (Other multi). D) Average relative abundance for the top 15 taxa across the populations sampled in each field for the cabbage stem flea beetle.

In contrast, symbionts of the cabbage stem flea beetle included a wide range of species. The 15 most abundant genera included several insect-associated microbes: *Aeromonas spp., Wolbachia spp., Pseudomonas spp*. and *Serratia spp*. Of these, *Wolbachia* spp. and *Pseudomonas* spp. dominated (Fig. 2D). Intra-seasonal variation was a key factor influencing the microbial community in the beetles, with populations surveyed in autumn having a different profile than summer populations (F_1,71_= 4.56; p = 0.033; Fig. 2D). The rarefaction curve is shown in Fig. S2.

Below, we describe the top-down and bottom-up effects for aphids and beetles separately.

### Landscape effects on natural enemies and aphids

We used piecewise structural equation models (pSEM) to identify the landscape processes influencing natural enemies, aphids, and aphid endosymbionts within cereal fields.

Only parasitoids, but not carabid natural enemies, responded significantly to landscape variables with seminatural habitat a key factor driving parasitoid abundance (Fig. 3A). Structural equation modelling identified configurational complexity (average field size) across the landscape as a significant factor influencing the abundance of the two spring/summer-abundant aphid species *S. avenae* and *M. dirhodum* (Fig. 3B) - fewer aphids were observed in landscapes with larger fields (Fig. S3). None of the three aphid species were directly influenced by host crop or seminatural habitat proportion in the landscape, but *M. dirhodum* were more abundant in sites with greater crop heterogeneity (Fig. 3A; Fig. S3). The proportion of seminatural habitat increased parasitoid wasp (Ichneumonidae and Braconidae) abundance, which in turn was linked to lower *M. dirhodum* abundance (Fig. 3A-B). Thus, landscape structure had both direct and indirect effects on spring/summer-abundant cereal aphids. This contrasted with the autumn-abundant aphid, *R. padi*, whose abundance was not associated with any landscape or enemy variable measured (Table S1; Table S2). Full model results and summary of all tested paths are detailed in Tables S1-4.

**Fig. 3.**
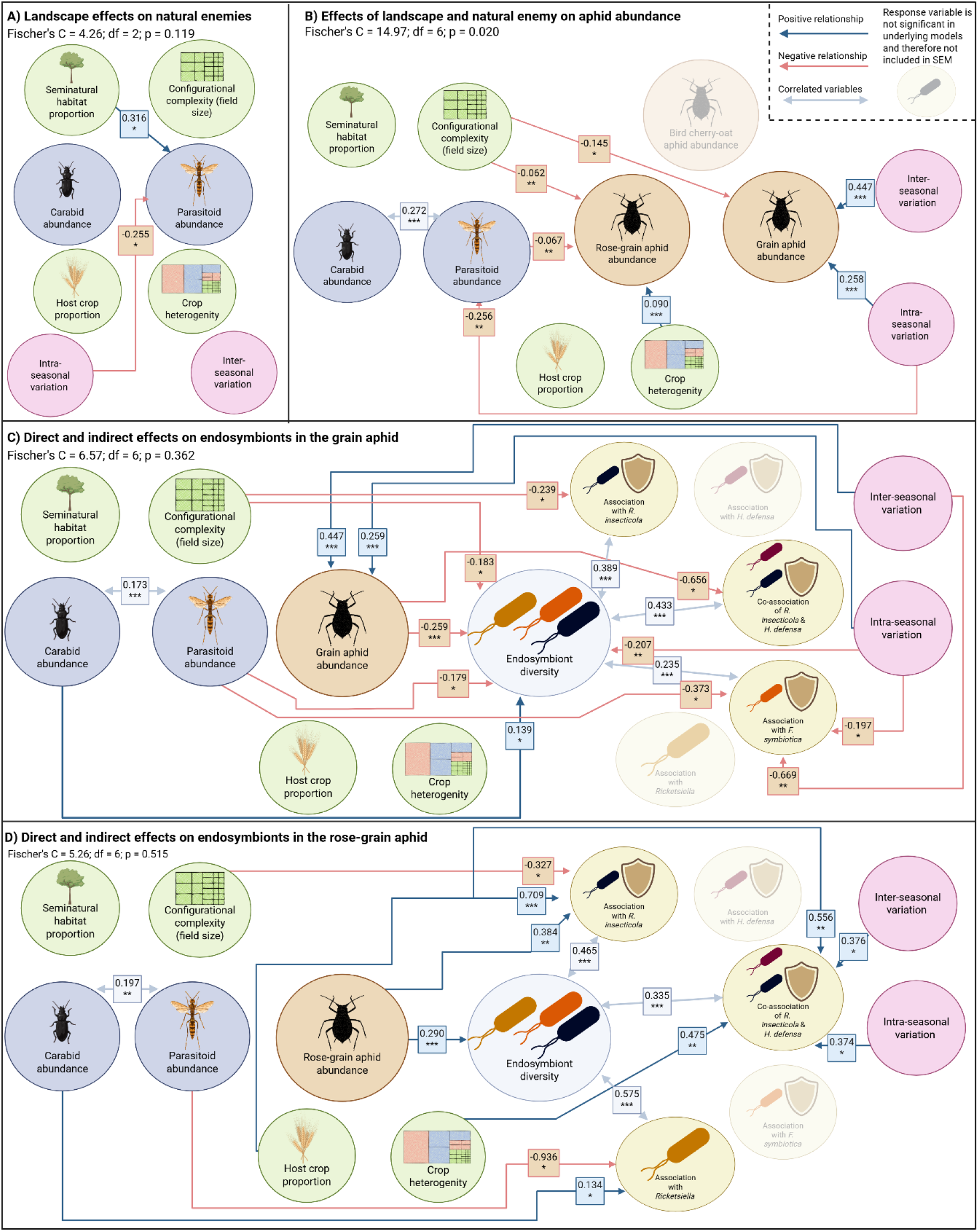
Piecewise Structural Equation Models (pSEM) of insect-landscape and aphid-endosymbiont interactions: A-B) Overall insect abundance pSEMs; C) Grain aphid, *S. avenae*, pSEM; D), Rose-grain aphid, *M. dirhodum*, pSEM. In B, C, and D natural enemies were included as explanatory variables to support model convergence, and correlations between the two natural enemy groups were included in these models. In all pSEMs, red paths (solid line) denote negative interactions while blue ones show positive relationships. To support visualisation, only the significant paths are shown; see Supplementary Tables S1-9 for all tested pathways and the full statistical results for each underlying model. Landscape variables are calculated based on the analysis of land use maps of 500m-radius circular areas around sampled fields. Dataset observations: 170 observations (natural enemy pSEM), 672 observations (aphid abundance pSEM), 432 observations (*S. avenae* endosymbiont pSEM), 221 observations (*M. dirhodum* endosymbionts pSEM). Asterisks denote levels of significance: * p < 0.05; ** p <0.01; *** p <0.001.

### Cascading effects of landscape regulation on aphid-endosymbiont associations

We used pSEM to examine the impact the landscape has on aphid-endosymbiont relationships for *S. avenae* and *M. dirhodum* (Fig. 3C-D); there were too few *R. padi* samples to build a reliable model. For *M. dirhodum*, aphids collected from fields with larger aphid populations hosted a more diverse cohort of endosymbionts (Fig. S4A). However, the opposite effect was observed for *S. avenae*, where aphids from larger populations hosted fewer endosymbionts (Fig. S4B). For *S. avenae*, this was associated with fewer aphids co-hosting *R. insecticola* and *H. defensa* (Fig. S4C). In addition, parasitoid abundance was also negatively related to endosymbiont diversity in *S. avenae*, though with contrasting effects between years (Fig. S4D).

In contrast, we observed a direct negative relationship between *M. dirhodum* abundance and parasitoid pressure (Fig. S4E). There was no significant association between parasitoid abundance and endosymbiont diversity in *M. dirhodum*. Yet, fewer *M. dirhodum* hosted *Rickettsiella* symbionts in areas with higher parasitoid abundance (Fig. S4F). Across both aphid species, we also observed significant associations between endosymbionts and carabid beetle abundance. A positive relationship was observed between *S. avenae* symbiont diversity and carabid abundance (Fig. S4G), while in *M. dirhodum* there was no direct effect on endosymbiont diversity, but an increase in *Ricketsiella spp*. association as carabid abundance increased (Fig. S4H).

### Bottom-up landscape processes influence aphid-microbe relationships

The pSEM also identified bottom-up processes directly linking landscape characteristics with endosymbiont communities (Fig. 3C-D). Across the two years of sampling, *S. avenae* aphids collected from landscapes with larger fields consistently showed lower endosymbiont diversity in individual aphids (Fig. 3C). Aphids sampled from these landscapes also showed a reduced tendency to host *R. insecticola* (Fig. S5A-B). For *M. dirhodum* we observed significant effects of crop heterogeneity, with aphids sampled from sites with a greater crop compositional heterogeneity more likely to be co-infected with *R. insecticola* and *H. defensa* (Fig. S5C). Similar associations were observed between increasing host crop proportion and greater association with *R. insecticola* in single (Fig. S5D) and co-infections with *H. defensa* (S5E) in *M. dirhodum*. Of these, only *R. insecticola* was influenced by total aphid abundance within the field (Fig. S5F), representing a potential indirect path back to the cascading effect of landscape-mediated regulation of aphid populations.

Inter- and intra-seasonal variation also influenced aphid and natural enemy abundances and aphid endosymbiont diversity (Fig. 3), indicating dynamic effects through the season and across years. Full aphid endosymbiont model results are detailed in Tables S5-9.

### Landscape configuration and host crop composition are key drivers of microbiome assembly in the cabbage stem flea beetle

Due to low goodness of fit of the assembled pSEM, landscape effects on the beetle microbiome were identified through individual mixed effects models. We assessed three microbiome community metrics: Relative abundance, microbiome diversity, microbiome dominance. Fig. 4 provides a summary of the key findings.

**Fig. 4.**
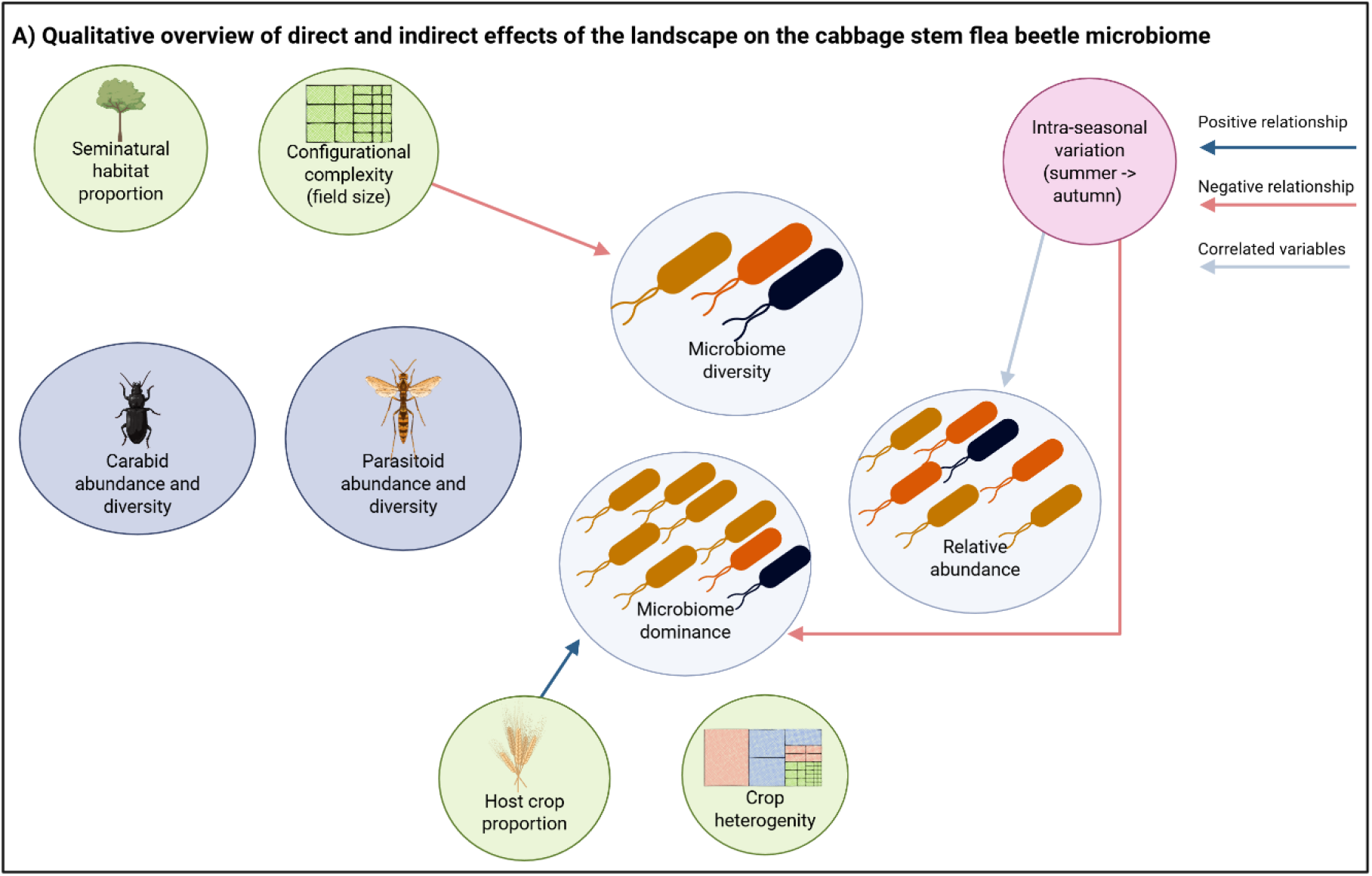
Graphical summary of the landscape parameters found to influence the microbial community in the cabbage stem flea beetle. Dataset comprises four individual beetles per survey site, with 72 beetle samples across all sites (full dataset) and 56 beetle samples in the autumn sites (autumn dataset). Results of mixed effects models linking the illustrated variables are shown in Tables S10, S11.

Microbiome diversity decreased as configurational complexity (field size) increased (X^2^_1_ = 4.25; p = 0.039; Fig. S6A). In contrast, microbiome dominance increased as host crop proportion increased in the landscape (X^2^_1_ = 8.28; p = 0.004; Fig. S6B). Microbiome dominance was also affected between the seasons with a greater dominance in summer populations (X^2^_1_ = 4.29; p = 0.038; Fig. S6B). Full model results are detailed in Table S10. These trends broadly follow our observations for cereal aphids (decreasing diversity as field sizes increase) and for *M. dirhodum* (host crop proportion impacts).

The natural enemy communities at the autumn sampling sites were previously characterised (19). Here, parasitoid and carabid pressure in the previous month and assessment month did not directly affect the microbiome metrics examined. Full model results are detailed in Table S11.

## Discussion

Many herbivorous insects form associations with microbial symbionts that confer a range of functional benefits, including protection from natural enemies, detoxification of plant secondary metabolites, and improved fitness [3, 14]. The prevalence and abundance of microbial symbionts can vary between insect populations [13, 15, 16], indicating that the insect-microbe relationship is influenced by external factors. Our results explicitly show for the first time that habitat configurational complexity (average field size) and the proportion of host-plant crop in the landscape, combined with natural enemy abundance, can act as direct and indirect drivers of microbial symbiosis in herbivorous insects.

### Configurational complexity and natural enemy pressure shape insect-symbiont relationships

The average size of agricultural fields is an effective measure of configurational complexity, with a landscape comprising multiple smaller fields indicative of greater connectivity [25]. Generally, configurational complexity has been found to influence insect populations by enhancing insect activity and dispersal: Greater connectivity of agricultural land (i.e., smaller average field size) promotes spillover and movement of insects between fields and habitats [25, 26]. Greater connectivity can also affect natural enemy activity, increasing top-down regulation of herbivorous insects [25].

Configurational complexity is known to influence cereal aphids [27], with increasing field size leading to smaller aphid populations [27, 28]. We found that configurational complexity influences the partnerships made between aphids and their endosymbionts, including increased association with the protective endosymbiont *R. insecticola*. Increased landscape connectivity (smaller fields in the landscape) can promote natural enemy activity, and therefore promote selection for more protective microbiomes; providing a direct mechanism for increased hosting of protective symbionts. This could have potential consequences on agricultural sustainability as aphids could persist in the landscape longer, with consequences of increased viral spread; particularly as protective endosymbionts can impact aphid interactions with host plants and plant viruses [29, 30].

We observed a negative relationship between parasitoid pressure and endosymbiont diversity in individual aphids (*S. avenae*), *F. symbiotica* association (*S. avenae*), and *Rickettsiella spp*., association (*M. dirhodum*). Although these endosymbionts are not experimentally documented to confer parasitoid resistance, our results suggest that these endosymbionts confer potentially costly phenotypes to the aphid when parasitoid pressure is high, and that less diverse microbiomes are beneficial in these cases [31]. Recent analysis across several aphid species identified a link between parasitoid attack networks and the distribution of defensive microbial partners within aphid populations [32], indicating that aphid species with a shared parasitoid network are likely to have comparable microbiomes. Furthermore, observations in the peach potato aphid, *Myzus persicae*, found that the presence of *Rickettsiella* increased parasitoid wasp attack rate [33], indicating that this microbial partner can confer a significant cost on the host aphid, particularly when parasitoids are present.

The protective phenotypes conferred by aphid endosymbionts (*H. defensa, R. insecticola*) can vary between aphid species, endosymbiont strain, and specific endosymbiont-aphid-parasitoid combinations [34, 35, 36], with protective phenotypes in *H. defensa* potentially mediated by the gene repertoire of toxin-encoding bacteriophages [37]. Recent research highlighted strain-level variation within protective microbes as a significant driver behind parasitoid protection [32, 34], therefore parasitoid pressure could be selecting for specific strains of protective microbes, with these shifts going undetected in our analysis. Comparable studies examining this relationship from the microbial perspective found similar trends, with the dominant parasitoid species shaping *H. defensa* strain-level variation [38]. Alternatively, *S. avenae* association with *H. defensa* and *R. inscticola* can cascade up the food chain, influencing parasitoid and hyperparasitoid communities [39]; thus the direct ecosystem-scale effect of these protective endosymbionts could act at the parasitoid level and not the aphid-endosymbiont level.

### Increased host crop availability supports more diverse microbiomes

In many chewing insects, microbial partners support the host insect by helping to detoxify or sequester potentially harmful plant secondary metabolites [14]. The cabbage stem flea beetle feeds primarily from the Brassicacae family, where highly toxic glucosinolates act as a central component of plant defence against herbivory and need to be detoxified during feeding (15). The core microbiome of various flea beetle species includes *Pantoea sp., Wolbachia sp*., and *Aeromonas sp*., where these core species are integral in facilitating herbivory, plant adaptation, and secondary metabolite detoxification [14, 40, 41].

Research in three different *Altica* flea beetle species found limited variation in α-diversity metrics between the microbiome of beetles sampled from different host crops, suggesting a core microbiome is required for successful herbivory [41]. This supports similar observations in the cabbage stem flea beetle where core microbial species support glucosinolate detoxification [14]. The cabbage stem beetles in our study hosted similar microbial species as core components of their microbiome, showing consistent associations across different systems. We further demonstrated that the overall dominance of these core species increased in tandem with host crop proportion in the landscape. This indicates that these beneficial species increase in relative abundance as insect exposure, and access to, the host plant increases. This is potentially driven by an increase in the requirement of microbiome-mediated detoxification processes, triggering a shift in the core microbiome as beetle populations are exposed to more host crop availability. In a parallel finding, host crop proportion was also highlighted as a landscape factor affecting microbial association in the rose-grain aphid, *M. dirhodum*. Here, the main effect was centred around increased association with the potentially protective microbial partner *R. insecticola* either alone or in co-association with *H. defensa*.

### Conclusion: Landscape drivers of insect-microbe relationships

Here, we use field-collected samples across a replicated gradient of agricultural landscapes to examine how landscape features and natural enemy pressure affect the formation of microbial partnerships in herbivorous insects. We find that the configuration of the landscape combined with the abundance of natural enemies are key factors driving endosymbiont association in both aphids and flea beetles, potentially acting via indirect and uncharacterised paths. Host crop proportion is also an important factor underpinning microbiome structure in a leaf-chewing beetle, potentially acting through selection for microbiomes better able to detoxify plant secondary metabolites. Understanding how insect symbionts can alter population dynamics of their hosts in the context of landscape configuration, is an important yet currently missing input for models predicting pest and pathogen spread in managed landscapes.

## Materials & Methods

### Survey sites and landscape characterisation

Field sites were located in a key area of arable production in Lower Saxony, Germany (Fig. S1). All fields were sown and managed by the host farmer using standard agronomic practices. Cereal sites were sown in autumn and surveyed the following summer during the period of peak aphid activity (June and July). Cereal sites were divided into 18 fields in 2021 and 17 in 2022. Eighteen rapeseed fields were visited in 2021, divided into four sites surveyed in summer (sown in 2020) and 14 sites surveyed in autumn (sown in 2021). Rapeseed surveys corresponded with the period of adult beetle emergence (summer) and migration of adults into host crops following summer aestivation (autumn). Host farmers were contacted prior to each survey to confirm that no insecticide had been recently applied to the fields.

The landscape around each site was mapped at a 500 m spatial scale, a scale sufficient to identify top-down and bottom-up effects on the focal insects [18, 42]. The landscape surrounding each field was characterised using open-access digital crop-cover maps. Detailed crop maps were obtained from the Lower Saxony federal database on agricultural development (Servicezentrum Landentwicklung und Agrarförderung). These geodata contain information on crop species grown in each field across Lower Saxony. We retrieved data on semi-natural habitat from the Lower Saxony ATKIS database (ATKIS-Objektartenkatalog). QGIS v.3.24.3 was used for landscape characterisation.

The following landscape variables were considered: 1) Host crop proportion; 2) crop heterogeneity; 3) seminatural habitat proportion; and 4) configurational complexity (average field size, Ha). Crop heterogeneity was calculated by extracting the total number of fields for each crop species present in the landscape and calculating Shannon’s diversity Index. Seminatural habitat was calculated by summing the total proportion for forest, woodland, heath, moor, swamp, and uncultivated land, as done previously [18].

For cereal sites, there were no consistent significant associations between landscape variables across the two years of data collection; although there were some correlations within specific years (Fig. S7). In 2021, host crop proportion was correlated with crop heterogeneity (r=-0.617), and configurational complexity with seminatural habitat proportion (r=0.625). In 2022, only crop heterogeneity and seminatural habitat proportion were correlated (r=-0.574). There was no correlation between surrounding landscape variables for the rapeseed sites (Fig. S8).

### Experimental design: Cereal fields

Each field comprised two 100 m long transects with five equidistantly spaced 1 m^2^ quadrats. Both transects ran parallel to the field edge, the first transect was 5 m from the field boundary and the second transect was 25 m from the field boundary; the only deviation for this was site 2021-27 where transect length was limited by field size, here transects were 70 m long with 14 m distances between quadrats. The same edge type (i.e. a crop– crop boundary) was used at each field site to reduce edge effects.

Each site was visited up to three times in summer (June – July) and aphid abundance was scored across each quadrat, divided into *S. avenae, M. dirhodum, R. padi*, and total abundance. The number of mummified aphids was also noted. In 2021, visitations were carried out w/c 7^th^ June, 21^st^ June, and 5^th^ July; two fields, 2021-35 and 2021-41, were only surveyed twice as the crop had reached maturity by the final survey date. In 2022 fields were visited w/c 6^th^ June and 20^th^ June. Where present, one sample for each of the following aphid categories were collected from each quadrat: *R. padi, M. dirhodum, S. avenae* – green clone, *S. avenae* pink/brown clone, and *S. avenae* black/purple clone. Up to three aphid mummies were collected per quadrat: *Praon spp, Aphidius spp*., and *Aphelinus spp*. Samples were stored at −20 °C until DNA extraction. During some visits very few aphids were found within the transects, on these occasions a crop walk was carried out to sample aphids, with a minimum distance of *c*. 20 m per sampling location.

A pair of insect sampling traps were installed in the central quadrat of each transect during each visit and left exposed for 48 h. Each trap pair comprised one yellow pan trap and one pitfall trap. Traps were 1/3 filled with water containing household washing detergent. At the request of host farmers, pan traps were placed on the soil surface, not elevated at canopy height. The contents from each trap type were pooled and traps were processed for each transect, with the Hymenoptera and Coleoptera identified to family level. The Stresemann taxonomic key was used to support arthropod identification [43]. Trap surveys at three sites (2022-78, 2022-79, 2022-82) differed slightly, here three trapping locations were used placed at equidistant from the field edge towards the field centre.

### Experimental design: Rapeseed fields

At summer sites, four pan traps were installed along the edge of the field with at least 20 m between trap locations. For autumn sites, pan traps were installed at the central location along two 100 m long transects, with one transect running along the field edge and a parallel transect 20 m into the crop. The same edge type was used at each field site to reduce edge effects. Pan traps were exposed for 24 h (summer sampling) or 48 h (autumn sampling). Beetles were stored in 96% ethanol in the freezer until DNA extraction, with four individual beetles processed per field.

### DNA extraction and microbiome characterisation

#### Aphids

The Chelex® extraction method was used to extract DNA from the aphid and mummy samples following the process described in [44]. Briefly, A 5% Chelex® 100-resin (Bio-Rad, Germany) solution was made and heated to 60°C. Then 10 µL Chelex® solution was mixed with 1.5 µL Proteinase K (20 mg L-1, Macherey-Nagel GmbH & Co. KG, Germany) and added to each sample. Aphids were homogenised using sterile pipette tips and mummies with a sterile micropestle. After homogenisation, a further 80 µL Chelex® solution was added, the sample was vortexed and incubated for 1.5 h at 56 °C with occasional mixing. After incubation, samples were centrifuged at max speed for 15 minutes and the solution was transferred to a clean Eppendorf tube, diluted 1:2 with TE buffer (10 mM Tris-HCl pH 7.5; 1 mM EDTA pH 8.0), and stored at −20 °C. An extraction blank was included with each batch of DNA extracts. Successful DNA extraction was confirmed using a PCR marker for the obligate endosymbiont *Buchnera aphidicola*; only successful extractions were processed further. Aphid endosymbionts was characterised using three multiplex PCR assays described in Beekman, *et al*. [8]. The Beekman, *et al*. [8] multiplex assays can detect the main endosymbionts of aphids, including *Spiroplasma spp., Regiella insecticola, Hamiltonella defensa, Rickettsiella sp., Fukatsuia symbiotica, Serratia symbiotica, Rickettsia spp*., and *Arsenophonus spp*.

PCR assays were conducted in a final reaction volume of 12 µL. PCR contents comprised 1 µL DNA and 6 µL 2X Kappa2G Fast PCR Ready Mix (Merck, Germany). Nuclease-free DEPC-treated water (CarlRoth, Germany) was used to make the final reaction volume to 12 µL. Positive DNA (mixed DNA containing positive extracts for all target microbial species) was included as a positive control and DNA-free PCR mastermix was included as PCR negative control. Positive reactions were detected by separation of PCR products on a 1% agarose gel stained with GelRed® (Biotium, Germany) and visualisation under UV light; a 100 bp DNA ladder (ThermoFischer, Germany) was used to estimate band size. All PCR assays were conducted in a Biometra TRIO 48, Thermocycler (Analytik Jena, Germany).

#### Beetles

Individual beetles were surface-sterilised in bleach, rinsed three times with d.H_2_O, further sterilised by soaking in 90% ethanol, and rinsed a further three times with d.H_2_O. DNA was extracted using the salting out protocol. Briefly, beetle tissue was homogenised using a micropestle in 500 µL TNS buffer (1 mM Tris-base, 1 mM NaCl, 2% SDS) with 12.5 µL Proteinase K. Samples were incubated at 56 °C for 3.5 h with occasional vortexing, heated to 96 °C for ten minutes, and treated with 20 µL RNAse A for ten minutes at 56 °C. Samples were mixed with 500 µL phenol:chloroform:isopropanol (25:24:1), incubated for five minutes at room temperature, mixed with 500 µL chloroform, incubated for an additional five minutes at room temperature, and centrifuged at maximum speed for 15 minutes. The upper aqueous layer was removed, mixed with 2.5x cold 96% ethanol and 5M NaCl, and incubated overnight in the freezer. Samples were centrifuged at full speed for 20 minutes to pellet DNA, washed with 70% ethanol and centrifuged at full speed for another 20 minutes. Pellets were washed a second time, air-dried under a laminar flow hood and resuspended in 85 µL TE buffer (10 mM Tris-Hcl pH 7.5, 1mM EDTA pH 8.0). DNA concentration (ng µL^−1^), volume, and quality were measured using an Eppendorf BioPhotomoter (Eppendorf, Germany) with a 1 mm excitation wavelength in an Eppendorf UVette (Eppendorf, Germany). Extracted DNA was diluted to *c*. 30 ng µL^−1^ and used as template in 16S sequencing on an Illumina MiSeq V3 platform at LGC Genomics GmBH (Berlin, Germany). Library preparation, QC, and sequencing was carried out by LGC Genomics GmBH. Primer sequences targeted the hypervariable region: 515FY (GTGYCAGCMGCCGCGGTAA) and 926Rjed (CCGYCAATTYMTTTRAGTTT). Samples were also prepared for fungal microbiome assessment through ITS sequencing, but very few fungal sequences were obtained. Therefore, analysis focussed on bacterial microbes.

Data pre-processing was carried out by LGC Genomics GmbH (Berlin, Germany). This involved demultiplexing reads using Illumina Bcl2fastq v2.20 software; up to two mismatches or Ns were allowed in the barcode read. Reads were sorted using the inline barcodes, one mismatch was allowed. After sorting the sequence was clipped; reads with missing barcodes, one-sided barcodes, or conflicting barcode pairs were discarded. The sequence adapter was clipped from all reads and reads with a final length <100 bases were discarded. Finally, primer sequences were identified and clipped from each read; up to three mismatches per primer pair were allowed. After data pre-processing the “primer-clipped” reads were used as input and the DADA^2^ approach [45] was followed to process the data. Reads were trimmed to 200 bases, merged, and used to construct amplicon sequence variant tables. Sequences were aligned to the SILVA rDNA 16S database (v.138.1). Plastid sequences were identified and removed and a subsample of extraction blanks were included to identify lab-introduced contaminants, the “isContaminant” function in the decontam package [46] was used to identify and remove potential contaminants. ASVs with a low read number (<1000 reads) were removed. ASVs were pre-processed using proportional normalisation, a normalisation method that has been found to work better when comparing among communities [47]. These data were used in the analysis below.

### Statistical analysis

Data were analysed in R Studio running R v.4.5.1. The following additional packages were used: tidyverse (v.2.0.0), ape (5.8-1), car (v.3.1-5), DHARMa (v.0.4.7), glmmTMB (v.1.1.14), vegan (v.2.7-3), lme4 (v.2.0-1), MuMIn (v.1.47.1), piecewiseSEM (v.2.3.1), ggplot2 (v.4.0.2), ggpubr (v.0.6.3).

In all models detailed below survey site was included as a random factor, accounting for random variation in agronomic practice. We used a mixture of negative binomial generalised linear mixed effects models (carabid abundance, parasitoid abundance, aphid abundance, and presence/absence of aphid endosymbionts) and linear mixed effects models (aphid endosymbiont diversity, beetle microbiome abundance, beetle microbiome diversity, beetle microbial dominance). In each model, we used a Variance Inflation Factor (VIF) cut-off value of five to define collinear variables [48] and removed any explanatory variables that breached this threshold. Models were tested for significance using analysis of deviance tests (Type II Wald X^2^ tests) and the fitted-residual plots of the models were assessed to check model suitability and conformance to model assumptions. In models where at least one bottom-up landscape feature was identified as a significant explanatory variable we used Moran’s I to test for spatial autocorrelation of the model residuals. Spatial autocorrelation was only detected in two models, the *M. dirhodum* and *R. padi* abundance models (Table S12).

#### Aphid and natural enemy abundance

The abundance of natural enemies (carabid and parasitoid wasps) and aphids was assessed at each site. Natural enemy abundance was pooled across the two trap types and examined in response to landscape variables, visit number, and assessment year. Aphid abundance was examined in three separate models (one per aphid species) in response to landscape variables, visit number, assessment year, total carabid abundance (summed across traps) and total parasitoid abundance (summed across traps).

To identify direct and indirect effects we built a two structural equation models (pSEMs), one for natural enemy abundance and one for aphid abundance. Each pSEM comprised all mixed effects models that contained at least one significant landscape variable, with the aphid pSEM accounting for correlation between carabid and parasitoid abundance. We extracted standardised coefficients and evaluated model fit using Fisher’s C statistic.

#### Aphid endosymbionts

For each individual aphid we scored presence or absence of any facultative microbial partner as a binary variable, summed the total number of microbial partners for each aphid, and calculated the microbiome diversity (Shannon’s diversity). In our initial analysis, we used the full dataset containing information from all three aphid species, with aphid species, survey year, visit number, and all landscape metrics included as explanatory variables. Upon identifying significant differences between the three aphid species, we subset the data into three additional datasets (one per aphid species), and used these datasets to conduct a more detailed analysis of bottom-up and top-down effects. Here, we examined presence or absence of any facultative endosymbiont, total number of microbial partners, Shannon’s diversity, and the presence or absence of key endosymbionts: *H. defensa, R. insecticola, F. symbiotica, Ricketsiella, and* co-infection of *R. insecticola* and *H. defensa*. Explanatory variables in these models included those detailed above alongside a covariate of total abundance of the focal aphid species (i.e., total *S. avenae* abundance observed in each field at each visitation in the *S. avenae* models), parasitoids, and carabids for each survey site.

Finally, to identify direct and indirect effects on aphid-symbiont relationships we built a single pSEM for each aphid species. Each pSEM contained all mixed effects models that comprised at least one significant landscape variable and was built to account for potential covariance between endosymbiont diversity and presence/absence of individual endosymbiont species. We extracted standardised coefficients and evaluated model fit using Fisher’s C statistic. Due to a low number of *R. padi* samples we were unable to develop a pSEM for *R. padi*.

#### Beetle microbiome

We calculated three key microbiome diversity indices (relative abundance, microbial diversity, microbial dominance) and examined these in response to landscape variables, season (summer vs autumn), natural enemy abundance in the preceding week, and natural enemy abundance in the assessment week. We explored pSEM approaches, but this was not appropriate due to unsatisfactory model convergence.

## Acknowledgements

The authors thank Petra Melloh, Alexander Manentzos, Eric Mühlnikel, Meike Meyer, Anna-Lena Heitkamp, Kristina Dauven, Christoph Harms, Ruth Simon, and David Schöner (Leibniz University Hannover, Germany) for their assistance with summer fieldwork and processing of insect traps. The authors also extend thanks to Dr Antonino Malacrinò (Clemson University, USA) for helpful discussions on 16S analysis.

