## Supplementary Material for "Top-down and bottom-up landscape processes influence the formation of microbial partnerships in herbivorous insects"

**Fig. S1:** Location of the study sites. Top: Cereal sites categorised into sampling year (2021, 2022). Bottom: Rapeseed sites categorised into sampling season (summer, autumn).

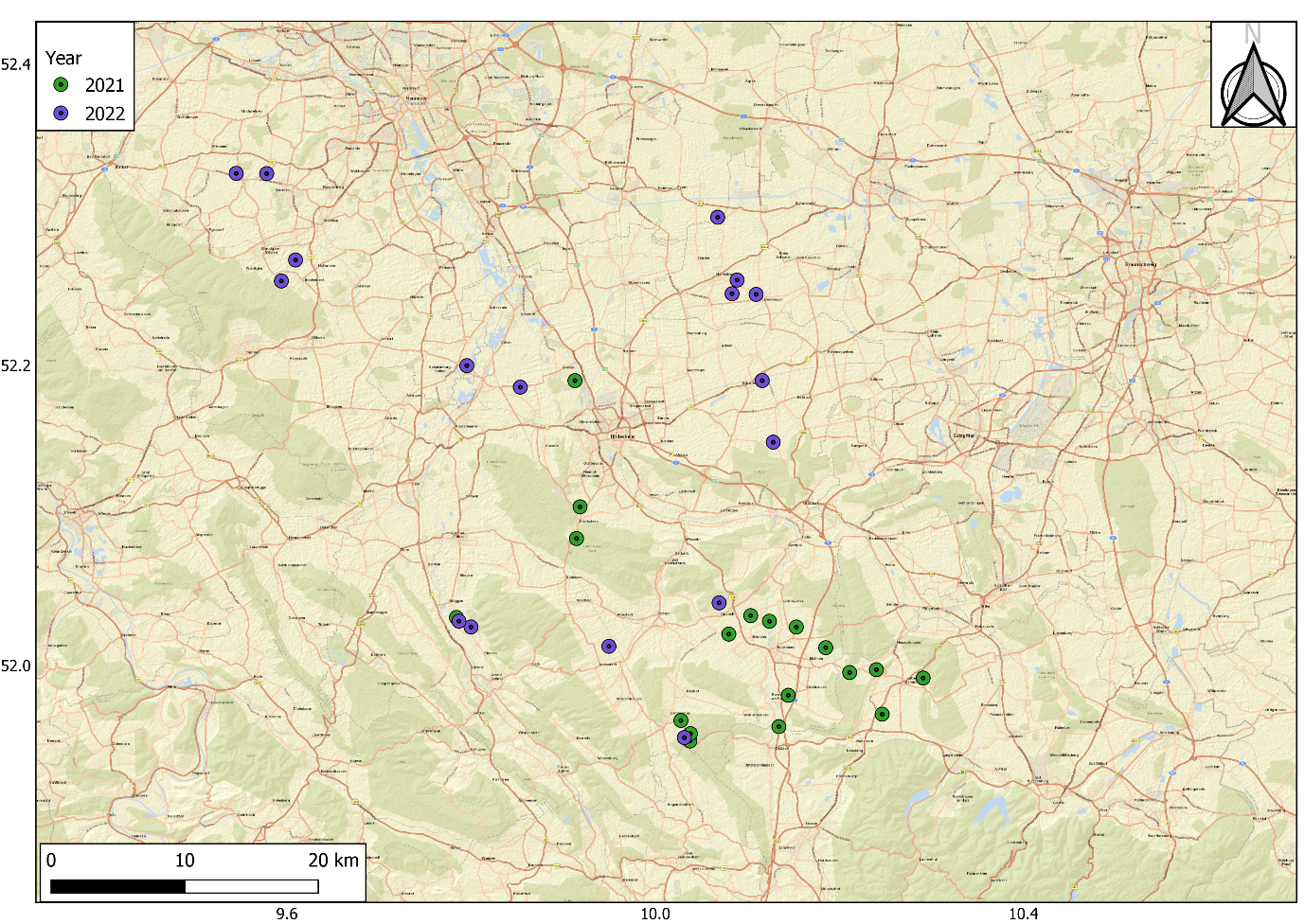

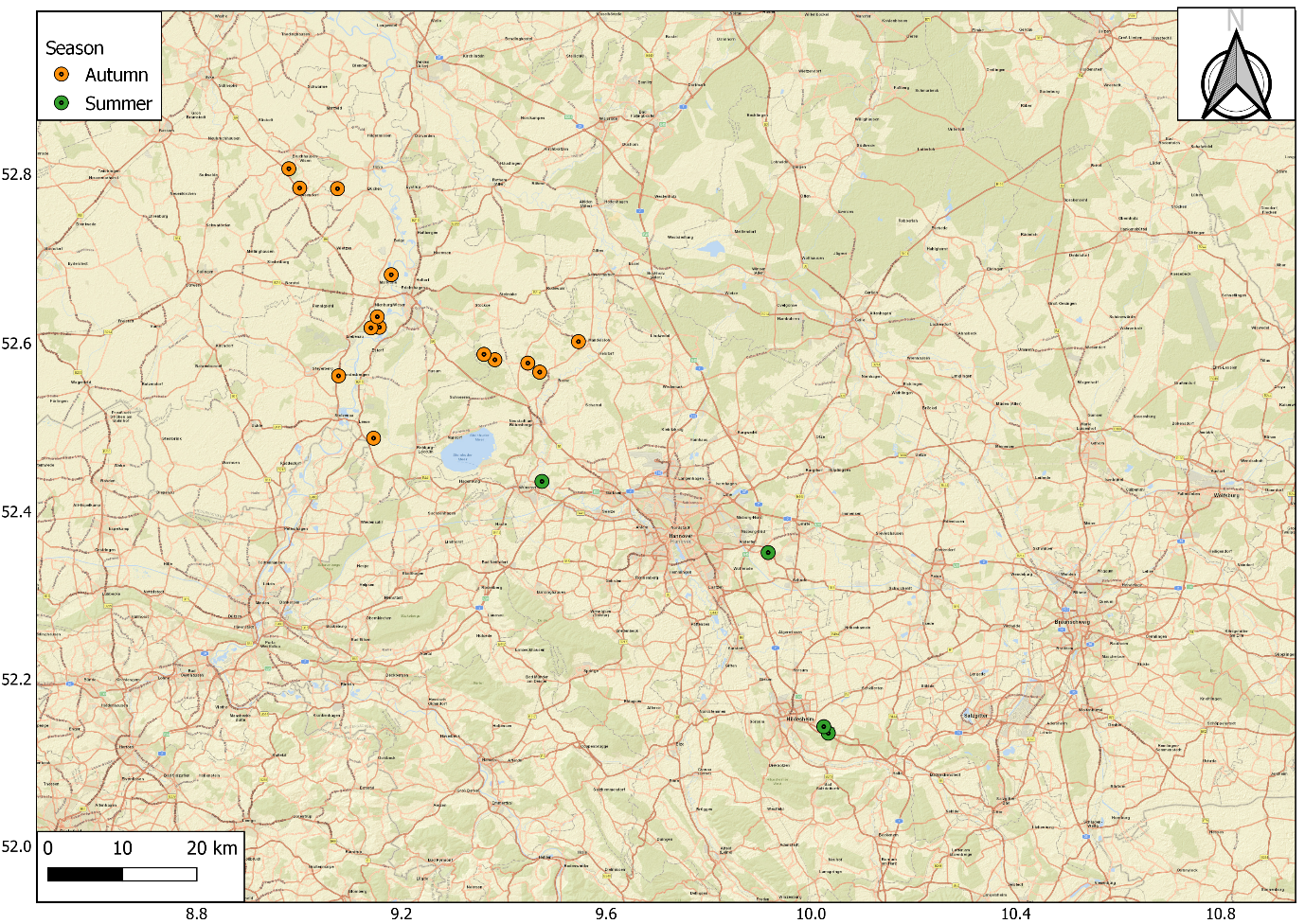

**Fig. S2:** Rarefaction curves for cabbage stem flea beetle 16S data.

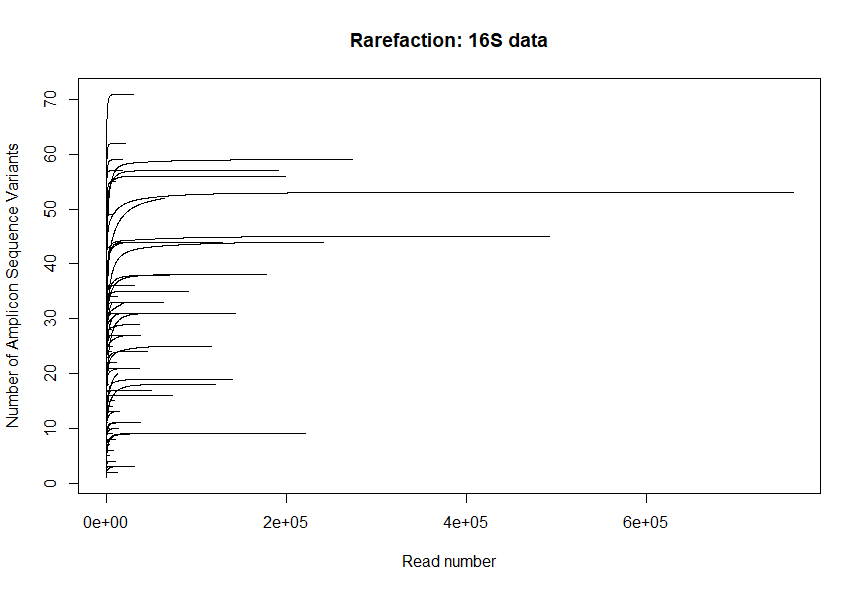

**Fig. S3:** Influence of bottom-up landscape factors on cereal aphid abundance. All landscape metrics are assessed over a 500 m radius. See Table S1 for associated statistical results.

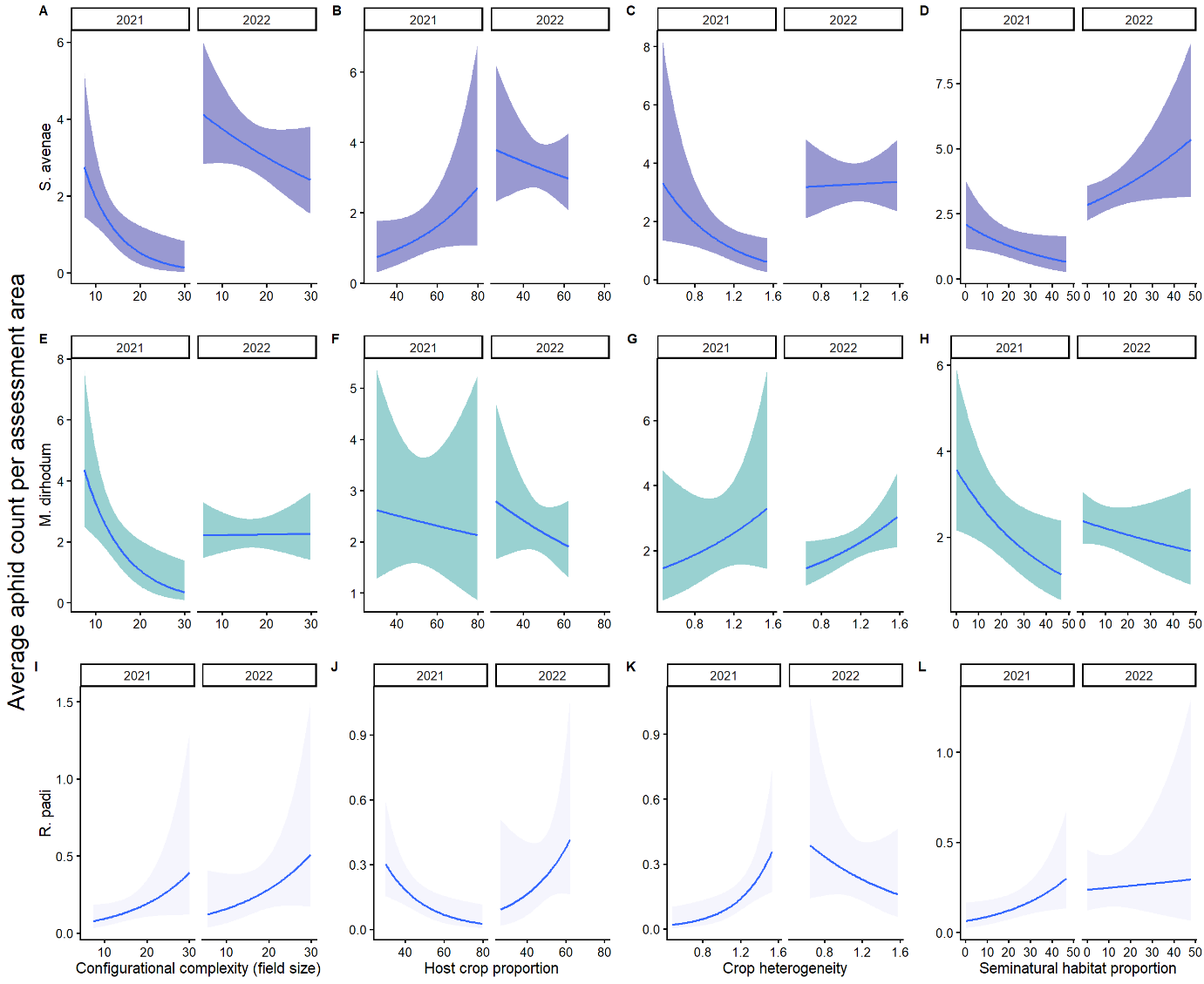

**Fig. S4:** Influence of top-down factors on endosymbiont partnerships in *S. avenae* (SA) and *M. dirhodum* (MD). Only significant relationships are displayed.

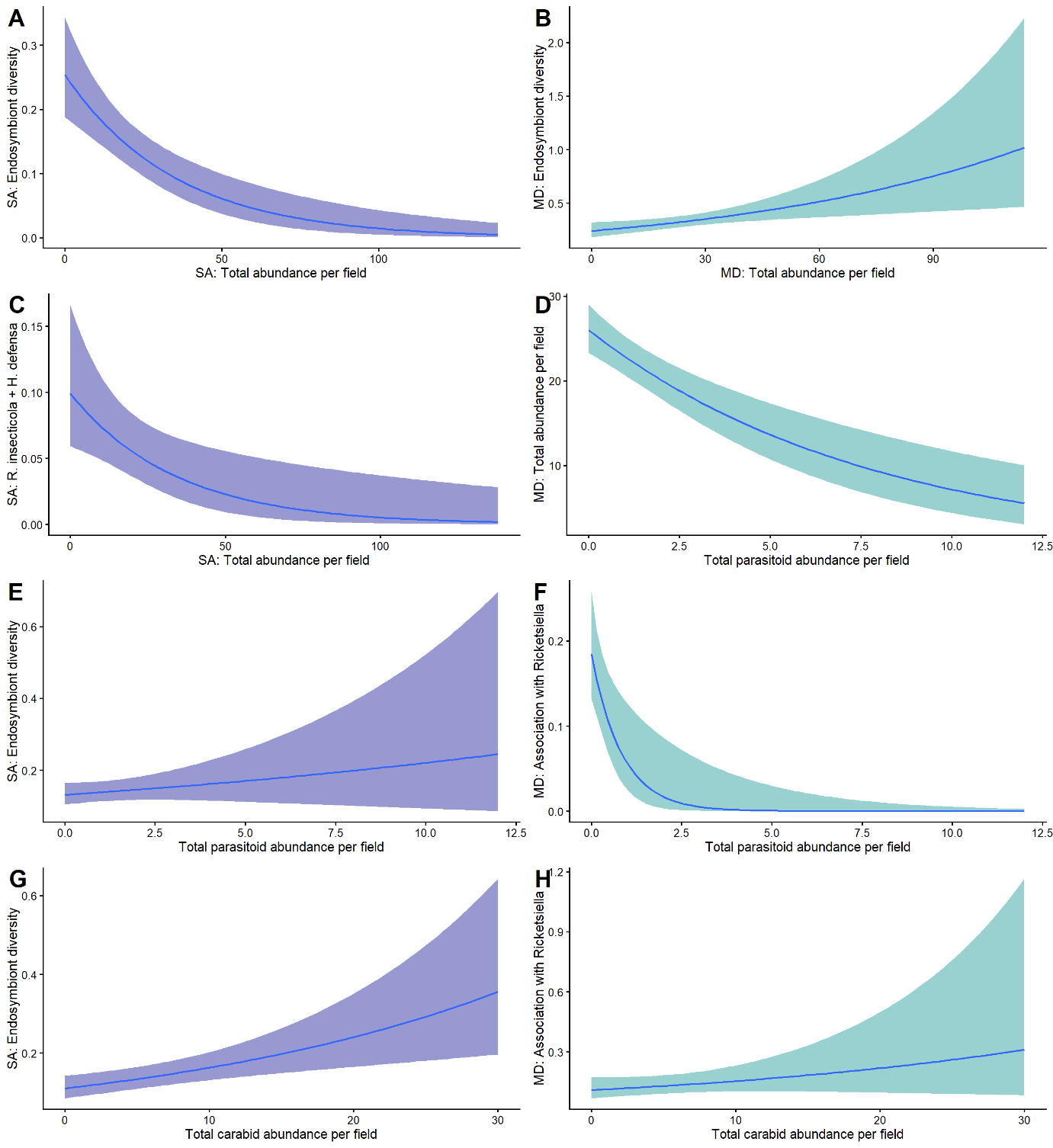

**Fig. S5:** Influence of bottom-up factors on microbial partnerships in *S. avenae* (SA) and *M. dirhodum* (MD). Only significant relationships are displayed. All landscape metrics are assessed over a 500 m radius.

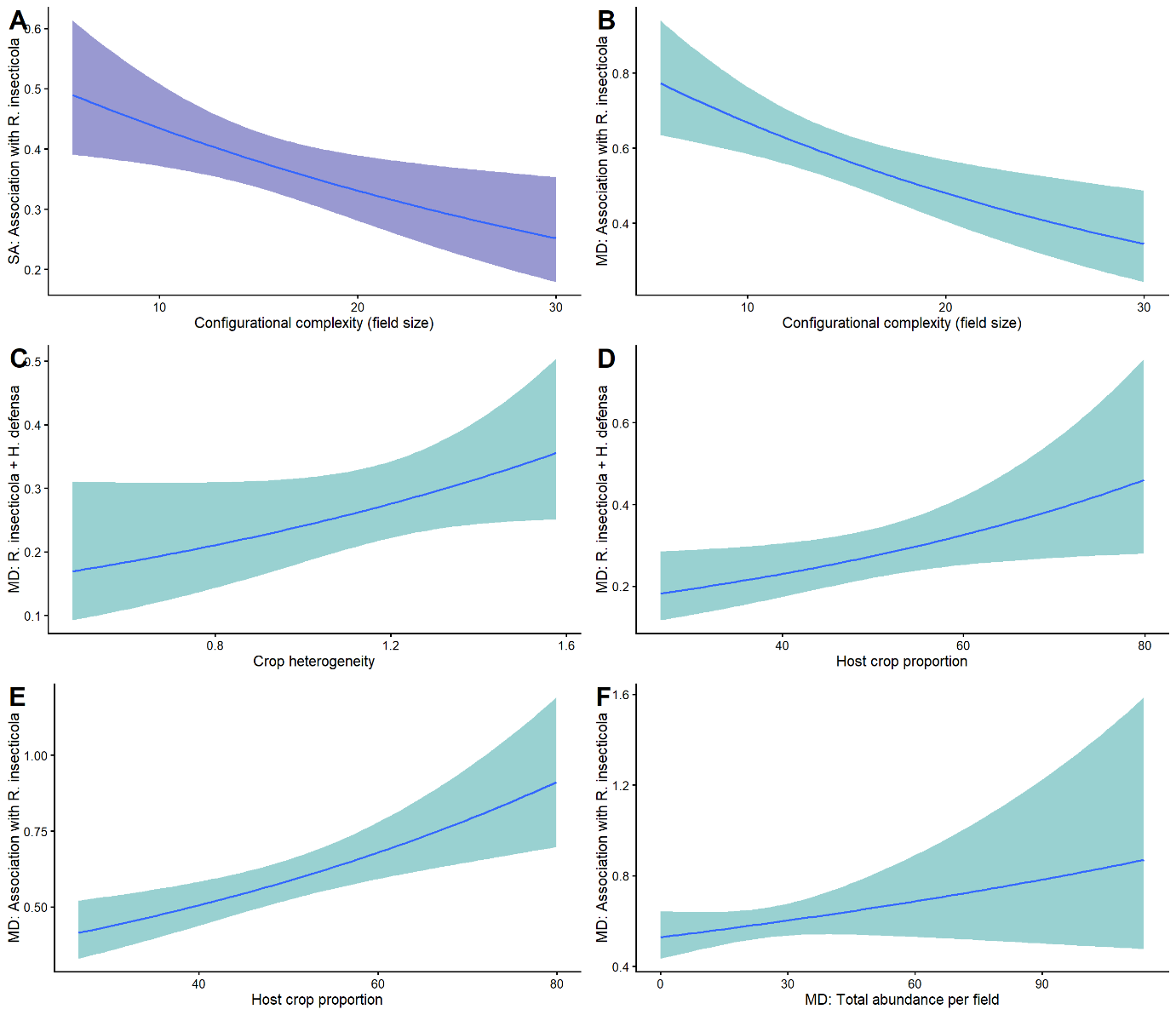

**Fig. S6:** Influence of bottom-up factors on the beetle microbiome. All landscape metrics are assessed over a 500 m radius.

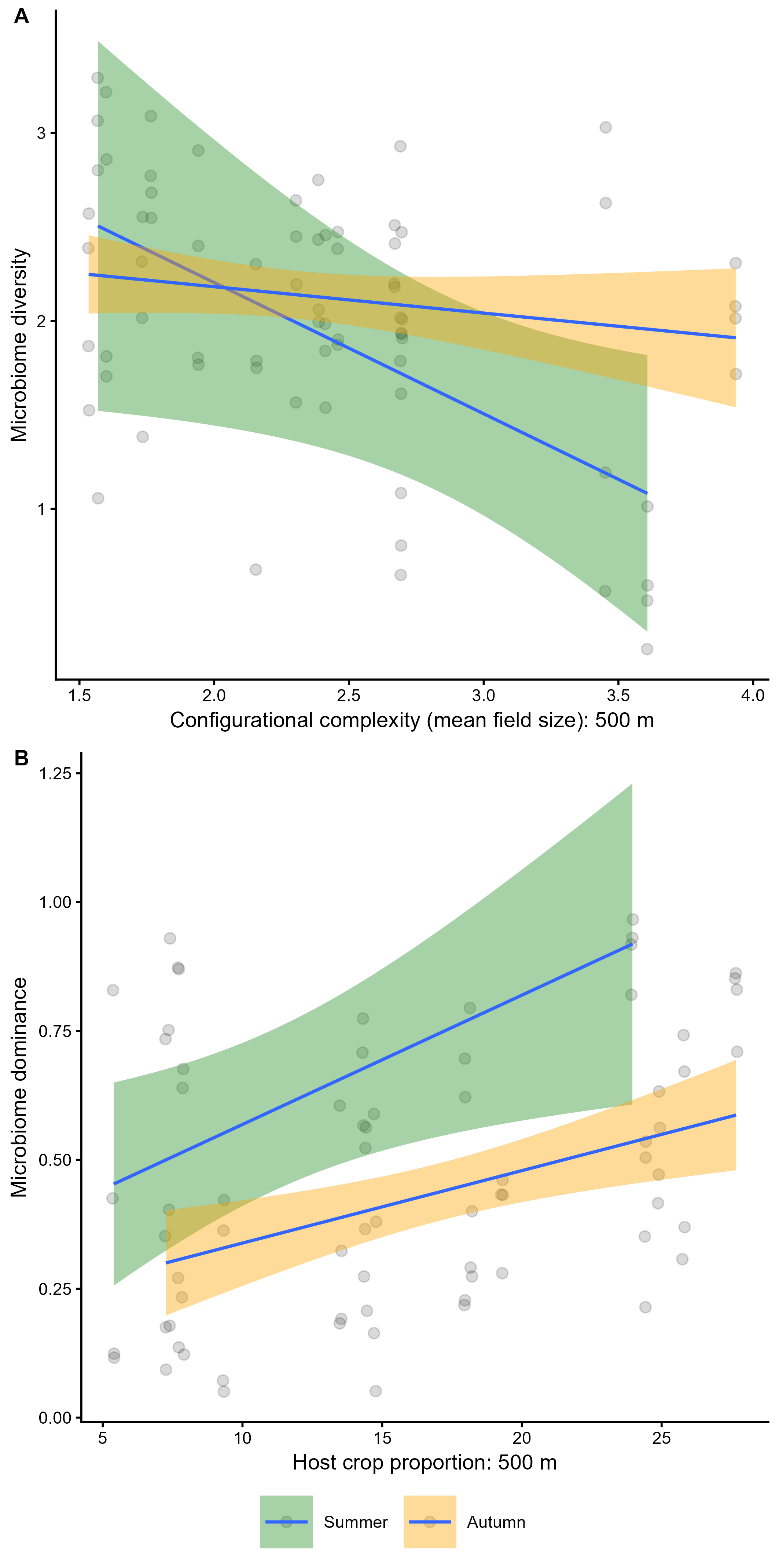

**Fig. S7:** Correlation between landscape variables for cereal sites across the two years.

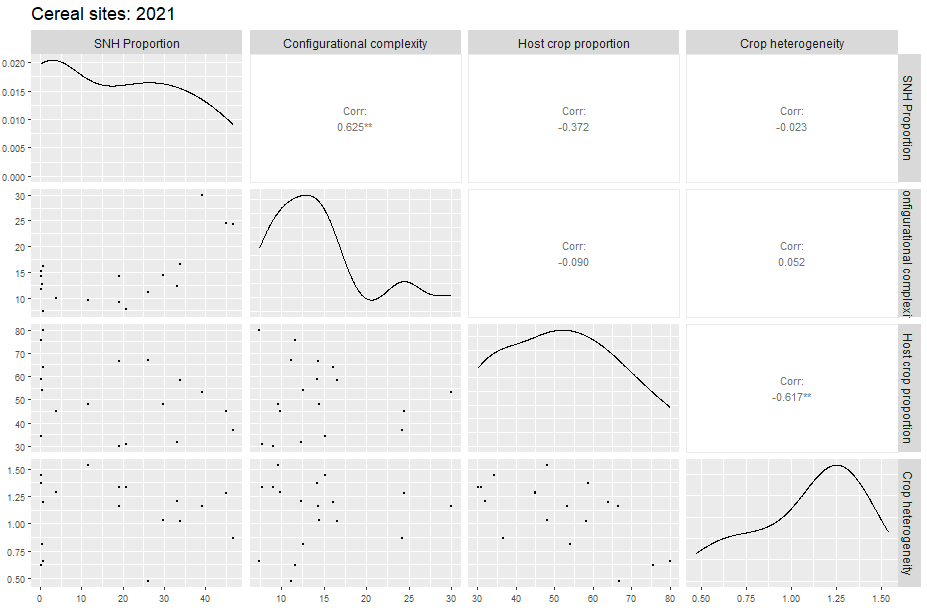

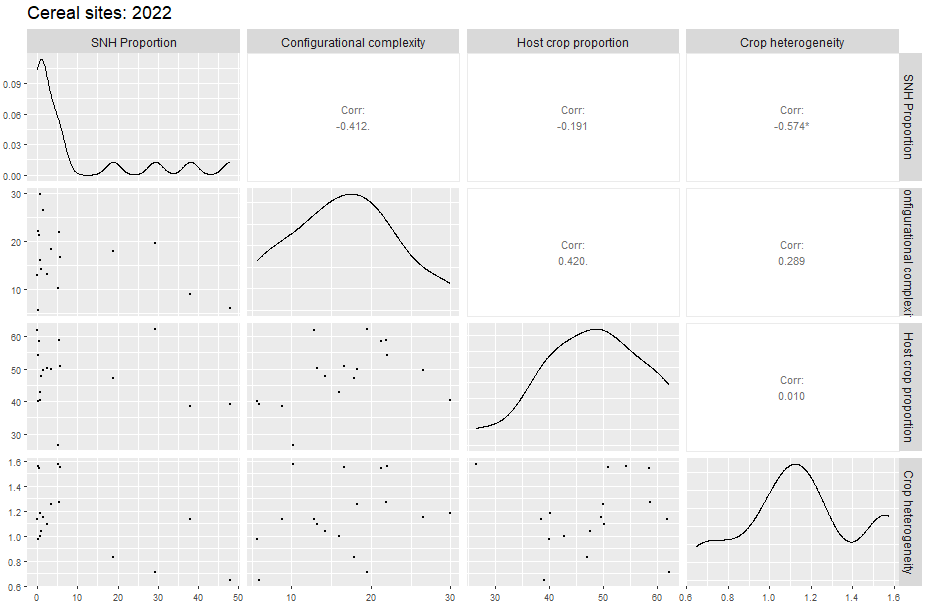

**Fig. S8:** Correlation between landscape variables for rapeseed sites.

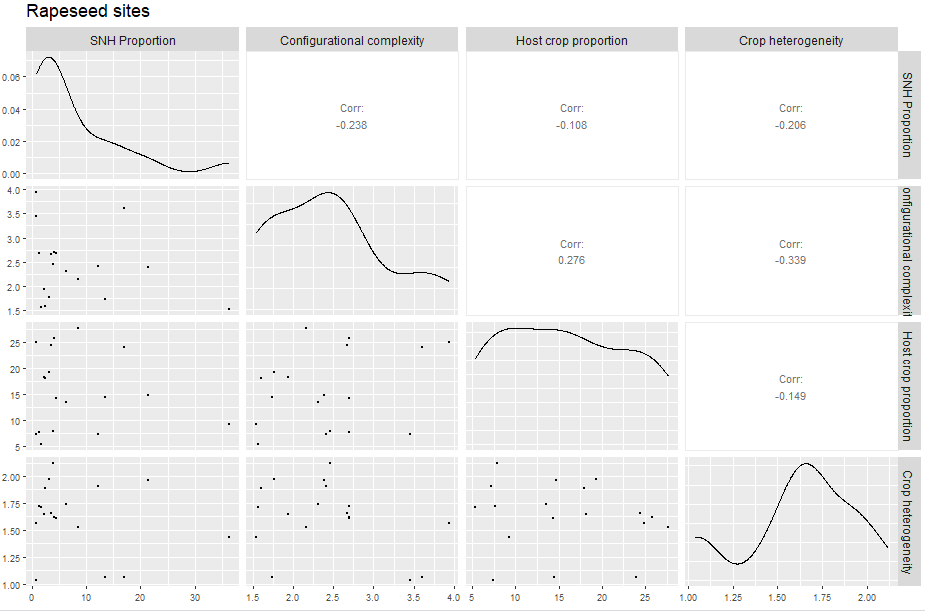

**Table S1:** Full results of all aphid abundance models.

| **Response variable** | **Χ^2^** | **DF** | **p-value** |
| --- | --- | --- | --- |
| *Sitobion avenae* abundance | | | |
| Crop heterogeneity | 0.09 | 1 | 0.763 |
| Field size | 4.11 | 1 | **0.043** |
| Host crop proportion | 1.14 | 1 | 0.291 |
| SNH proportion | 0.00 | 1 | 0.965 |
| Carabid abundance | 2.13 | 1 | 0.144 |
| Parasitoid abundance | 2.14 | 1 | 0.143 |
| Visit number | 17.13 | 1 | **<0.001** |
| Year | 27.47 | 1 | **<0.001** |
| *Metapolophium dirhodum* abundance | | | |
| Crop heterogeneity | 10.78 | 1 | **0.001** |
| Field size | 6.74 | 1 | **0.009** |
| Host crop proportion | 0.06 | 1 | 0.801 |
| SNH proportion | 0.57 | 1 | 0.451 |
| Carabid abundance | 0.11 | 1 | 0.742 |
| Parasitoid abundance | 7.07 | 1 | **0.007** |
| Visit number | 3.19 | 1 | 0.074 |
| Year | 1.96 | 1 | 0.161 |
| *Rhopalosiphum padi* abundance | | | |
| Crop heterogeneity | 0.15 | 1 | 0.694 |
| Field size | 3.22 | 1 | 0.072 |
| Host crop proportion | 1.38 | 1 | 0.229 |
| SNH proportion | 0.41 | 1 | 0.524 |
| Carabid abundance | 0.55 | 1 | 0.461 |
| Parasitoid abundance | 0.01 | 1 | 0.936 |
| Visit number | 0.98 | 1 | 0.321 |
| Year | 1.83 | 1 | 0.176 |

**Table S2:** Full results of all SEM paths for aphid abundance measurements.

| **Response** | **Predictor** | **Estimate** | **Std.Error** | **DF** | **Crit.Value** | **P.Value** | **Std.Estimate** |
| --- | --- | --- | --- | --- | --- | --- | --- |
| Sitobion | CropHet_500m | -0.2176 | 0.7246 | 642 | -0.3004 | 0.7639 | -0.0241 |
| Sitobion | Mean_Size_Ha_500m | -0.0603 | 0.0297 | 642 | -2.0283 | **0.0425** | -0.1415 |
| Sitobion | Prop_Cereal_500m | 0.0175 | 0.0166 | 642 | 1.0557 | 0.2911 | 0.0842 |
| Sitobion | SNH_Prop_500m | -6.00E-04 | 0.0133 | 642 | -0.0444 | 0.9645 | -0.0036 |
| Sitobion | Carabidae | 0.0301 | 0.0206 | 642 | 1.4608 | 0.1441 | 0.0722 |
| Sitobion | ParaAbundance | 0.1541 | 0.1053 | 642 | 1.4633 | 0.1434 | 0.0794 |
| Sitobion | Visit_Number_B | 0.9819 | 0.2372 | 642 | 4.1391 | **<0.001** | 0.2583 |
| Sitobion | Yearb | 2.3667 | 0.4515 | 642 | 5.2416 | **<0.001** | 0.4471 |
| Metapolophum | CropHet_500m | 1.3159 | 0.4007 | 642 | 3.2837 | **0.001** | 0.0904 |
| Metapolophum | Mean_Size_Ha_500m | -0.0423 | 0.0163 | 642 | -2.5961 | **0.0094** | -0.0616 |
| Metapolophum | Prop_Cereal_500m | 0.0023 | 0.009 | 642 | 0.2512 | 0.8016 | 0.0067 |
| Metapolophum | SNH_Prop_500m | -0.0055 | 0.0073 | 642 | -0.7531 | 0.4514 | -0.0207 |
| Metapolophum | Carabidae | -0.0051 | 0.0155 | 642 | -0.3296 | 0.7417 | -0.0076 |
| Metapolophum | ParaAbundance | -0.2109 | 0.0793 | 642 | -2.6598 | **0.0078** | -0.0674 |
| Metapolophum | Visit_Number_B | -0.3596 | 0.2012 | 642 | -1.7873 | 0.0739 | -0.0587 |
| Metapolophum | Yearb | -0.4108 | 0.2933 | 642 | -1.4006 | 0.1613 | -0.0482 |
| Rhopalosiphum | CropHet_500m | 0.5249 | 1.3382 | 642 | 0.3922 | 0.6949 | 0.0313 |
| Rhopalosiphum | Mean_Size_Ha_500m | 0.0936 | 0.0522 | 642 | 1.7931 | 0.073 | 0.1185 |
| Rhopalosiphum | Prop_Cereal_500m | -0.0357 | 0.0303 | 642 | -1.176 | 0.2396 | -0.0925 |
| Rhopalosiphum | SNH_Prop_500m | 0.015 | 0.0235 | 642 | 0.6365 | 0.5244 | 0.0488 |
| Rhopalosiphum | Carabidae | -0.0343 | 0.0465 | 642 | -0.738 | 0.4605 | -0.0444 |
| Rhopalosiphum | ParaAbundance | 0.0184 | 0.198 | 642 | 0.0931 | 0.9258 | 0.0051 |
| ~~Carabidae | ~~ParaAbundance | 0.2719 | - | 640 | 7.1492 | 0 | 0.2719 |

**Table S3:** Full results of all natural enemy abundance models.

| **Response variable** | **Χ^2^** | **DF** | **p-value** |
| --- | --- | --- | --- |
| Carabid abundance | | | |
| Crop heterogeneity | 0.18 | 1 | 0.668 |
| Field size | 0.01 | 1 | 0.911 |
| Host crop proportion | 0.56 | 1 | 0.454 |
| SNH proportion | 0.14 | 1 | 0.708 |
| Visit number | 1.69 | 1 | 0.193 |
| Year | 2.31 | 1 | 0.128 |
| Parasitoid abundance | | | |
| Crop heterogeneity | 2.35 | 1 | 0.125 |
| Field size | 0.14 | 1 | 0.717 |
| Host crop proportion | 0.27 | 1 | 0.600 |
| SNH proportion | 5.57 | 1 | **0.018** |
| Visit number | 6.31 | 1 | **0.012** |
| Year | 0.02 | 1 | 0.903 |

**Table S4:** Full results of all SEM paths for natural enemy abundance measurements.

| **Response** | **Predictor** | **Estimate** | **Std.Error** | **DF** | **Crit.Value** | **P.Value** | **Std.Estimate** |
| --- | --- | --- | --- | --- | --- | --- | --- |
| ParaAbundance | CropHet_500m | 1.0684 | 0.6958 | 124 | 1.5354 | 0.1247 | 0.196 |
| ParaAbundance | Mean_Size_Ha_500m | -0.011 | 0.0297 | 124 | -0.3695 | 0.7117 | -0.0413 |
| ParaAbundance | Prop_Cereal_500m | 0.008 | 0.0152 | 124 | 0.5242 | 0.6001 | 0.0655 |
| ParaAbundance | SNH_Prop_500m | 0.0316 | 0.0134 | 124 | 2.3609 | **0.0182** | 0.3161 |
| ParaAbundance | Visit_Number_B | -0.5713 | 0.2274 | 124 | -2.5126 | **0.012** | -0.2558 |
| ParaAbundance | Yearb | 0.0535 | 0.4375 | 124 | 0.1224 | 0.9026 | 0.0164 |
| Carabidae | CropHet_500m | -0.2693 | 0.6275 | 168 | -0.4292 | 0.6677 | -0.0845 |
| Carabidae | Mean_Size_Ha_500m | 0.0028 | 0.0248 | 168 | 0.1116 | 0.9111 | 0.0178 |
| Carabidae | Prop_Cereal_500m | -0.0107 | 0.0144 | 168 | -0.7482 | 0.4543 | -0.1508 |
| Carabidae | SNH_Prop_500m | -0.0042 | 0.0112 | 168 | -0.375 | 0.7077 | -0.0718 |
| Carabidae | Visit_Number_B | -0.195 | 0.1496 | 168 | -1.303 | 0.1926 | -0.1494 |
| Carabidae | Yearb | -0.4904 | 0.3225 | 168 | -1.5205 | 0.1284 | -0.2567 |

**Table S5:** Full results of endosymbiont models across full dataset (all aphid species combined).

| **Response variable** | **Χ^2^** | **DF** | **p-value** |
| --- | --- | --- | --- |
| Facultative endosymbiont presence | | | |
| Aphid species | 11.81 | 2 | **0.003** |
| Crop heterogeneity | 0.05 | 1 | 0.823 |
| Field size | 1.81 | 1 | 0.178 |
| Host crop proportion | 0.01 | 1 | 0.919 |
| SNH proportion | 0.01 | 1 | 0.941 |
| Visit number | 0.07 | 1 | 0.974 |
| Year | 0.02 | 1 | 0.890 |
| Total number of endosymbionts | | | |
| Aphid species | 41.63 | 2 | **<0.001** |
| Crop heterogeneity | 0.08 | 1 | 0.769 |
| Field size | 3.75 | 1 | 0.053 |
| Host crop proportion | 0.02 | 1 | 0.866 |
| SNH proportion | 0.25 | 1 | 0.617 |
| Visit number | 0.03 | 1 | 0.871 |
| Year | 3.54 | 1 | 0.059 |
| Facultative endosymbiont diversity | | | |
| Aphid species | 56.59 | 2 | **<0.001** |
| Crop heterogeneity | 0.23 | 1 | 0.631 |
| Field size | 4.04 | 1 | **0.044** |
| Host crop proportion | 0.01 | 1 | 0.926 |
| SNH proportion | 0.82 | 1 | 0.366 |
| Visit number | 0.95 | 1 | 0.328 |
| Year | 7.78 | 1 | **0.005** |

**Table S6:** Full results of all *S. avenae* endosymbiont models.

| **Response variable** | **Χ^2^** | **DF** | **p-value** |
| --- | --- | --- | --- |
| Facultative endosymbiont diversity | | | |
| Crop heterogeneity | 0.32 | 1 | 0.573 |
| Field size | 4.22 | 1 | **0.040** |
| Host crop proportion | 0.57 | 1 | 0.452 |
| SNH proportion | 0.12 | 1 | 0.733 |
| Year | 1.22 | 1 | 0.268 |
| Visit number | 9.32 | 1 | **0.002** |
| *S. avenae* abundance | 12.22 | 1 | **<0.001** |
| Carabid abundance | 5.80 | 1 | **0.016** |
| Parasitoid abundance | 7.69 | 1 | **0.005** |
| Association with *H. defensa* | | | |
| Crop heterogeneity | 1.38 | 1 | 0.240 |
| Field size | 0.70 | 1 | 0.402 |
| Host crop proportion | 0.62 | 1 | 0.432 |
| SNH proportion | 0.17 | 1 | 0.677 |
| Year | 1.45 | 1 | 0.229 |
| Visit number | 0.00 | 1 | 0.972 |
| *S. avenae* abundance | 0.98 | 1 | 0.321 |
| Carabid abundance | 0.30 | 1 | 0.582 |
| Parasitoid abundance | 0.12 | 1 | 0.729 |
| Association with *R. insecticola* | | | |
| Crop heterogeneity | 0.56 | 1 | 0.454 |
| Field size | 6.21 | 1 | **0.013** |
| Host crop proportion | 0.00 | 1 | 0.973 |
| SNH proportion | 1.08 | 1 | 0.299 |
| Year | 0.19 | 1 | 0.663 |
| Visit number | 0.56 | 1 | 0.354 |
| *S. avenae* abundance | 0.65 | 1 | 0.419 |
| Carabid abundance | 0.85 | 1 | 0.356 |
| Parasitoid abundance | 0.57 | 1 | 0.451 |
| Association with *F. symbiotica* | | | |
| Crop heterogeneity | 0.739 | 1 | 0.389 |
| Field size | 0.329 | 1 | 0.566 |
| Host crop proportion | 3.23 | 1 | 0.072 |
| SNH proportion | 0.34 | 1 | 0.561 |
| Year | 8.89 | 1 | **0.003** |
| Visit number | 5.16 | 1 | **0.023** |
| *S. avenae* abundance | 3.00 | 1 | 0.083 |
| Carabid abundance | 3.12 | 1 | 0.077 |
| Parasitoid abundance | 8.76 | 1 | **0.003** |
| Association with *Ricketsiella spp.* | | | |
| Crop heterogeneity | 0.097 | 1 | 0.756 |
| Field size | 0.43 | 1 | 0.511 |
| Host crop proportion | 0.00 | 1 | 0.963 |
| SNH proportion | 0.01 | 1 | 0.929 |
| Year | 0.11 | 1 | 0.743 |
| Visit number | 3.49 | 1 | 0.064 |
| *S. avenae* abundance | 0.73 | 1 | 0.391 |
| Carabid abundance | 0.11 | 1 | 0.736 |
| Parasitoid abundance | 0.03 | 1 | 0.870 |
| Co-association: *R. insecticola* and *H. defensa* | | | |
| Crop heterogeneity | 0.03 | 1 | 0.847 |
| Field size | 0.10 | 1 | 0.748 |
| Host crop proportion | 0.99 | 1 | 0.320 |
| SNH proportion | 0.15 | 1 | 0.699 |
| Year | 2.47 | 1 | 0.116 |
| Visit number | 0.34 | 1 | 0.557 |
| *S. avenae* abundance | 6.00 | 1 | **0.014** |
| Carabid abundance | 0.10 | 1 | 0.752 |
| Parasitoid abundance | 0.42 | 1 | 0.518 |

**Table S7:** Full results of all SEM paths for *S.avenae* endosymbiont associations.

| **Response** | **Predictor** | **Estimate** | **Std.Error** | **DF** | **Crit.Value** | **P.Value** | **Std.Estimate** |
| --- | --- | --- | --- | --- | --- | --- | --- |
| FacEndoDiv | CropHet_500m | 0.0577 | 0.1024 | 25.8883 | 0.316 | 0.5789 | 0.0585 |
| FacEndoDiv | Mean_Size_Ha_500m | -0.009 | 0.0044 | 31.8393 | 4.173 | **0.0494** | -0.1825 |
| FacEndoDiv | Prop_Cereal_500m | -0.0018 | 0.0023 | 25.6129 | 0.5638 | 0.4596 | -0.0693 |
| FacEndoDiv | SNH_Prop_500m | -7.00E-04 | 0.0019 | 30.7094 | 0.1154 | 0.7364 | -0.0331 |
| FacEndoDiv | Yearb | -0.0697 | 0.0629 | 45.9363 | 1.2092 | 0.2772 | -0.1125 |
| FacEndoDiv | Visit_Numberb | -0.0949 | 0.0311 | 303.4764 | 9.1181 | **0.0027** | -0.2067 |
| FacEndoDiv | Sitobion | -0.0023 | 7e-04 | 182.9498 | 11.7471 | **8e-04** | -0.2598 |
| FacEndoDiv | Carabidae | 0.0061 | 0.0025 | 211.5356 | 5.6126 | **0.0187** | 0.1395 |
| FacEndoDiv | ParaAbundance | -0.0287 | 0.0104 | 247.752 | 7.4921 | **0.0066** | -0.1792 |
| R_insecticola | CropHet_500m | 0.4994 | 0.6672 | 432 | 0.7485 | 0.4541 | 0.078 |
| R_insecticola | Mean_Size_Ha_500m | -0.076 | 0.0305 | 432 | -2.4918 | **0.0127** | -0.2387 |
| R_insecticola | Prop_Cereal_500m | 5e-04 | 0.0153 | 432 | 0.0335 | 0.9733 | 0.0031 |
| R_insecticola | SNH_Prop_500m | -0.0136 | 0.0131 | 432 | -1.0386 | 0.299 | -0.105 |
| R_insecticola | Yearb | -0.1896 | 0.4352 | 432 | -0.4358 | 0.663 | -0.0472 |
| R_insecticola | Visit_Numberb | -0.2298 | 0.2482 | 432 | -0.9257 | 0.3546 | -0.0771 |
| R_insecticola | Sitobion | -0.004 | 0.005 | 432 | -0.8087 | 0.4187 | -0.0707 |
| R_insecticola | Carabidae | -0.0191 | 0.0207 | 432 | -0.9231 | 0.356 | -0.0674 |
| R_insecticola | ParaAbundance | -0.0652 | 0.0864 | 432 | -0.7546 | 0.4505 | -0.0627 |
| F_symbiotica | CropHet_500m | -3.3486 | 3.895 | 432 | -0.8597 | 0.3899 | -0.209 |
| F_symbiotica | Mean_Size_Ha_500m | -0.0733 | 0.1278 | 432 | -0.5734 | 0.5664 | -0.0919 |
| F_symbiotica | Prop_Cereal_500m | -0.1524 | 0.0848 | 432 | -1.7968 | 0.0724 | -0.3681 |
| F_symbiotica | SNH_Prop_500m | -0.033 | 0.0567 | 432 | -0.5815 | 0.5609 | -0.1017 |
| F_symbiotica | Yearb | -6.7376 | 2.2586 | 432 | -2.9831 | **0.0029** | -0.6699 |
| F_symbiotica | Visit_Numberb | -1.4729 | 0.6484 | 432 | -2.2717 | **0.0231** | -0.1975 |
| F_symbiotica | Sitobion | -0.0472 | 0.0272 | 432 | -1.733 | 0.0831 | -0.3311 |
| F_symbiotica | Carabidae | 0.1198 | 0.0678 | 432 | 1.7666 | 0.0773 | 0.169 |
| F_symbiotica | ParaAbundance | -0.9707 | 0.3279 | 432 | -2.9601 | **0.0031** | -0.373 |
| Ri_Hd | CropHet_500m | 0.2814 | 1.4544 | 432 | 0.1935 | 0.8466 | 0.0327 |
| Ri_Hd | Mean_Size_Ha_500m | -0.0162 | 0.0503 | 432 | -0.3216 | 0.7478 | -0.0378 |
| Ri_Hd | Prop_Cereal_500m | -0.0325 | 0.0326 | 432 | -0.9942 | 0.3201 | -0.146 |
| Ri_Hd | SNH_Prop_500m | -0.0108 | 0.0278 | 432 | -0.387 | 0.6987 | -0.0618 |
| Ri_Hd | Yearb | 1.648 | 1.0488 | 432 | 1.5713 | 0.1161 | 0.3052 |
| Ri_Hd | Visit_Numberb | -0.3519 | 0.5994 | 432 | -0.5871 | 0.5571 | -0.0879 |
| Ri_Hd | Sitobion | -0.0502 | 0.0205 | 432 | -2.4506 | **0.0143** | -0.6568 |
| Ri_Hd | Carabidae | -0.0192 | 0.0608 | 432 | -0.3156 | 0.7523 | -0.0505 |
| Ri_Hd | ParaAbundance | -0.1439 | 0.2228 | 432 | -0.6456 | 0.5185 | -0.103 |
| ~~Carabidae | ~~ParaAbundance | 0.173 | - | 430 | 3.6427 | 3e-04 | 0.173 |
| ~~R_insecticola | ~~FacEndoDiv | 0.3887 | - | 432 | 8.7379 | **0** | 0.3887 |
| ~~F_symbiotica | ~~FacEndoDiv | 0.2345 | - | 432 | 4.997 | **0** | 0.2345 |
| ~~Ri_Hd | ~~FacEndoDiv | 0.4329 | - | 432 | 9.9468 | **0** | 0.4329 |

**Table S8:** Full results of all *M. dirhodum* endosymbiont models.

| **Response variable** | **Χ^2^** | **DF** | **p-value** |
| --- | --- | --- | --- |
| Facultative endosymbiont diversity | | | |
| Crop heterogeneity | 0.65 | 1 | 0.420 |
| Field size | 2.89 | 1 | 0.089 |
| Host crop proportion | 4.02 | 1 | **0.045** |
| SNH proportion | 1.86 | 1 | 0.172 |
| Year | 1.81 | 1 | 0.179 |
| Visit number | 0.00 | 1 | 0.952 |
| *M. dirhodum* abundance | 12.92 | 1 | **<0.001** |
| Carabid abundance | 0.01 | 1 | 0.895 |
| Parasitoid abundance | 0.76 | 1 | 0.382 |
| Association with *H. defensa* | | | |
| Crop heterogeneity | 0.00 | 1 | 0.986 |
| Field size | 0.59 | 1 | 0.441 |
| Host crop proportion | 3.32 | 1 | 0.068 |
| SNH proportion | 0.43 | 1 | 0.511 |
| Year | 5.12 | 1 | **0.023** |
| Visit number | 0.43 | 1 | 0.513 |
| *M. dirhodum* abundance | 3.08 | 1 | 0.079 |
| Carabid abundance | 1.28 | 1 | 0.257 |
| Parasitoid abundance | 1.33 | 1 | 0.248 |
| Association with *R. insecticola* | | | |
| Crop heterogeneity | 2.74 | 1 | 0.097 |
| Field size | 4.29 | 1 | **0.038** |
| Host crop proportion | 11.08 | 1 | **<0.001** |
| SNH proportion | 2.45 | 1 | 0.118 |
| Year | 1.12 | 1 | 0.289 |
| Visit number | 1.41 | 1 | 0.235 |
| *M. dirhodum* abundance | 7.93 | 1 | **0.005** |
| Carabid abundance | 0.45 | 1 | 0.503 |
| Parasitoid abundance | 0.06 | 1 | 0.800 |
| Association with *F. symbiotica* | | | |
| Crop heterogeneity | 0.16 | 1 | 0.692 |
| Field size | 0.42 | 1 | 0.517 |
| Host crop proportion | 0.11 | 1 | 0.738 |
| SNH proportion | 0.06 | 1 | 0.804 |
| Year | 1.69 | 1 | 0.193 |
| Visit number | 1.52 | 1 | 0.218 |
| *M. dirhodum* abundance | 0.96 | 1 | 0.326 |
| Carabid abundance | 0.16 | 1 | 0.689 |
| Parasitoid abundance | 0.70 | 1 | 0.403 |
| Association with *Ricketsiella spp.* | | | |
| Crop heterogeneity | 0.93 | 1 | 0.334 |
| Field size | 0.17 | 1 | 0.684 |
| Host crop proportion | 0.49 | 1 | 0.481 |
| SNH proportion | 0.29 | 1 | 0.591 |
| Year | 0.19 | 1 | 0.661 |
| Visit number | 3.74 | 1 | 0.053 |
| *M. dirhodum* abundance | 0.36 | 1 | 0.551 |
| Carabid abundance | 3.95 | 1 | **0.047** |
| Parasitoid abundance | 4.32 | 1 | **0.038** |
| Co-association: *R. insecticola* and *H. defensa* | | | |
| Crop heterogeneity | 10.67 | 1 | **0.001** |
| Field size | 2.19 | 1 | 0.139 |
| Host crop proportion | 13.84 | 1 | **<0.001** |
| SNH proportion | 2.97 | 1 | 0.085 |
| Year | 4.48 | 1 | **0.034** |
| Visit number | 4.34 | 1 | **0.037** |
| *M. dirhodum* abundance | 1.87 | 1 | 0.172 |
| Carabid abundance | 1.35 | 1 | 0.246 |
| Parasitoid abundance | 0.02 | 1 | 0.887 |

**Table S9:** Full results of all SEM paths for *M. dirhodum* endosymbiont associations.

| **Response** | **Predictor** | **Estimate** | **Std.Error** | **DF** | **Crit.Value** | **P.Value** | **Std.Estimate** |
| --- | --- | --- | --- | --- | --- | --- | --- |
| FacEndoDiv | CropHet_500m | 0.1489 | 0.1847 | 25.5444 | 0.6402 | 0.431 | 0.1158 |
| FacEndoDiv | Mean_Size_Ha_500m | -0.0127 | 0.0074 | 27.1569 | 2.843 | 0.1032 | -0.1931 |
| FacEndoDiv | Prop_Cereal_500m | 0.0083 | 0.0041 | 23.7428 | 3.9652 | 0.0581 | 0.2861 |
| FacEndoDiv | SNH_Prop_500m | 0.0046 | 0.0033 | 26.0015 | 1.8341 | 0.1873 | 0.1741 |
| FacEndoDiv | Yearb | 0.152 | 0.1131 | 51.4114 | 1.7876 | 0.1871 | 0.1835 |
| FacEndoDiv | Visit_Numberb | 0.0037 | 0.0606 | 151.1006 | 0.0036 | 0.9523 | 0.0069 |
| FacEndoDiv | Metapolophum | 0.0069 | 0.0019 | 184.9741 | 12.5203 | **5.00E-04** | 0.2903 |
| FacEndoDiv | Carabidae | -7.00E-04 | 0.0052 | 189.4957 | 0.0168 | 0.8969 | -0.0096 |
| FacEndoDiv | ParaAbundance | -0.0151 | 0.0172 | 173.2512 | 0.7429 | 0.3899 | -0.077 |
| R_insecticola | CropHet_500m | 2.5274 | 1.5272 | 221 | 1.6549 | 0.0979 | 0.3174 |
| R_insecticola | Mean_Size_Ha_500m | -0.1326 | 0.064 | 221 | -2.0727 | **0.0382** | -0.3265 |
| R_insecticola | Prop_Cereal_500m | 0.126 | 0.0379 | 221 | 3.3295 | **9.00E-04** | 0.7033 |
| R_insecticola | SNH_Prop_500m | 0.0437 | 0.0279 | 221 | 1.5651 | 0.1176 | 0.269 |
| R_insecticola | Yearb | 0.9561 | 0.9019 | 221 | 1.0601 | 0.2891 | 0.1864 |
| R_insecticola | Visit_Numberb | 0.55 | 0.4628 | 221 | 1.1883 | 0.2347 | 0.1671 |
| R_insecticola | Metapolophum | 0.0568 | 0.0202 | 221 | 2.8152 | **0.0049** | 0.3844 |
| R_insecticola | Carabidae | 0.0272 | 0.0407 | 221 | 0.6692 | 0.5034 | 0.0615 |
| R_insecticola | ParaAbundance | -0.0423 | 0.1672 | 221 | -0.2529 | 0.8004 | -0.0349 |
| Rickettsiella | CropHet_500m | -2.0765 | 2.1485 | 221 | -0.9665 | 0.3338 | -0.1167 |
| Rickettsiella | Mean_Size_Ha_500m | 0.0304 | 0.0745 | 221 | 0.4075 | 0.6836 | 0.0335 |
| Rickettsiella | Prop_Cereal_500m | -0.0356 | 0.0505 | 221 | -0.704 | 0.4815 | -0.0888 |
| Rickettsiella | SNH_Prop_500m | 0.019 | 0.0354 | 221 | 0.5377 | 0.5908 | 0.0524 |
| Rickettsiella | Yearb | -0.5176 | 1.1806 | 221 | -0.4384 | 0.6611 | -0.0451 |
| Rickettsiella | Visit_Numberb | -1.3182 | 0.6814 | 221 | -1.9344 | 0.0531 | -0.1792 |
| Rickettsiella | Metapolophum | 0.0148 | 0.0247 | 221 | 0.5969 | 0.5505 | 0.0447 |
| Rickettsiella | Carabidae | 0.1327 | 0.0668 | 221 | 1.9869 | **0.0469** | 0.1342 |
| Rickettsiella | ParaAbundance | -2.5376 | 1.2204 | 221 | -2.0793 | **0.0376** | -0.9364 |
| Ri_Hd | CropHet_500m | 3.4371 | 1.052 | 221 | 3.2673 | **0.0011** | 0.4753 |
| Ri_Hd | Mean_Size_Ha_500m | -0.0735 | 0.0497 | 221 | -1.4802 | 0.1388 | -0.1993 |
| Ri_Hd | Prop_Cereal_500m | 0.0905 | 0.0243 | 221 | 3.7207 | **2.00E-04** | 0.5563 |
| Ri_Hd | SNH_Prop_500m | 0.0346 | 0.0201 | 221 | 1.7223 | 0.085 | 0.2347 |
| Ri_Hd | Yearb | 1.7549 | 0.829 | 221 | 2.117 | **0.0343** | 0.3767 |
| Ri_Hd | Visit_Numberb | 1.1172 | 0.5361 | 221 | 2.0839 | **0.0372** | 0.3737 |
| Ri_Hd | Metapolophum | 0.0206 | 0.0151 | 221 | 1.367 | 0.1716 | 0.1533 |
| Ri_Hd | Carabidae | -0.0582 | 0.0502 | 221 | -1.161 | 0.2456 | -0.145 |
| Ri_Hd | ParaAbundance | -0.0279 | 0.1961 | 221 | -0.1424 | 0.8867 | -0.0254 |
| ~~Carabidae | ~~ParaAbundance | 0.1977 | - | 219 | 2.9842 | 0.0032 | 0.1977 |
| ~~R_insecticola | ~~FacEndoDiv | 0.4653 | - | 221 | 7.7612 | 0 | 0.4653 |
| ~~Rickettsiella | ~~FacEndoDiv | 0.3345 | - | 221 | 5.2408 | 0 | 0.3345 |
| ~~Ri_Hd | ~~FacEndoDiv | 0.5754 | - | 221 | 10.3888 | 0 | 0.5754 |

**Table S10:** Full results for all cabbage stem flea beetle microbiome models: Bottom-up effects

| **Response variable** | **Χ^2^** | **DF** | **p-value** |
| --- | --- | --- | --- |
| Microbiome diversity | | | |
| Crop heterogeneity | 0.44 | 1 | 0.503 |
| Field size | 4.25 | 1 | **0.039** |
| Host crop proportion | 0.78 | 1 | 0.376 |
| SNH proportion | 2.88 | 1 | 0.089 |
| Season | 2.35 | 1 | 0.124 |
| Microbiome dominance | | | |
| Crop heterogeneity | 0.49 | 1 | 0.482 |
| Field size | 0.49 | 1 | 0.483 |
| Host crop proportion | 8.21 | 1 | **0.004** |
| SNH proportion | 0.16 | 1 | 0.686 |
| Season | 4.29 | 1 | **0.038** |
| Relative abundance | | | |
| Crop heterogeneity | 0.15 | 1 | 0.689 |
| Field size | 1.28 | 1 | 0.256 |
| Host crop proportion | 1.21 | 1 | 0.270 |
| SNH proportion | 2.64 | 1 | 0.104 |
| Season | 4.56 | 1 | **0.033** |

**Table S11:** Full results for all cabbage stem flea beetle microbiome models: Top-down effects, autumn samples only

| **Response variable** | **Χ^2^** | **DF** | **p-value** |
| --- | --- | --- | --- |
| Microbiome diversity | | | |
| Putative natural enemy diversity: September (previous month) | 0.27 | 1 | 0.607 |
| Parasitoid wasp abundance: September (previous month) | 0.16 | 1 | 0.686 |
| Putative natural enemy diversity: October (same date as beetle sampling) | 0.06 | 1 | 0.810 |
| Parasitoid wasp abundance: October (same date as beetle sampling) | 0.01 | 1 | 0.933 |
| Microbiome dominance | | | |
| Putative natural enemy diversity: September (previous month) | 0.12 | 1 | 0.725 |
| Parasitoid wasp abundance: September (previous month) | 1.10 | 1 | 0.294 |
| Putative natural enemy diversity: October (same date as beetle sampling) | 0.27 | 1 | 0.603 |
| Parasitoid wasp abundance: October (same date as beetle sampling) | 0.31 | 1 | 0.580 |
| Relative abundance | | | |
| Putative natural enemy diversity: September (previous month) | 0.04 | 1 | 0.847 |
| Parasitoid wasp abundance: September (previous month) | 0.22 | 1 | 0.638 |
| Putative natural enemy diversity: October (same date as beetle sampling) | 0.02 | 1 | 0.893 |
| Parasitoid wasp abundance: October (same date as beetle sampling) | 0.02 | 1 | 0.883 |

**Table S12:** Test of spatial autocorrelation on model residuals. Moran’s I test results for all models that contain a significant landscape variable.

| **Model** | **Observed** | **Expected** | **Sd** | **p-value** |
| --- | --- | --- | --- | --- |
| **Aphid and natural enemy abundance models** | | | | |
| Carabid abundance | -0.004 | -0.005 | 0.004 | 0.693 |
| Parasitoid abundance | -0.005 | -0.008 | 0.005 | 0.682 |
| *Sitobion avenae* abundance | -0.002 | -0.001 | 0.001 | 0.609 |
| *Metopolophium dirhodum* abundance | -0.005 | -0.001 | 0.001 | **<0.001** |
| *Rhopalosiphum padi* abundance | -0.003 | -0.001 | 0.001 | **0.042** |
| **Aphid microbiome: All species** | | | | |
| Association with any endosymbiont | -0.000 | -0.001 | 0.000 | 0.323 |
| Total number of endosymbionts | -0.000 | -0.001 | 0.000 | 0.184 |
| Endosymbiont diversity | -0.000 | -0.001 | 0.000 | 0.171 |
| **Aphid microbiome: *S. avenae* models** | | | | |
| Endosymbiont diversity | -0.000 | -0.002 | 0.001 | 0.174 |
| *Hamiltonella defensa* | -0.001 | -0.002 | 0.001 | 0.417 |
| *Regiella insecticola* | -0.001 | -0.002 | 0.001 | 0.689 |
| *Fukatsuia symbiotica* | -0.000 | -0.002 | 0.001 | 0.211 |
| *Ricketsiella spp.* | -0.000 | -0.002 | 0.001 | 0.217 |
| Co-infection: *H. defensa* + *R. insecticola* | -0.000 | -0.002 | 0.001 | 0.279 |
| **Aphid microbiome: *M. dirhodum* models** | | | | |
| Endosymbiont diversity | -0.000 | -0.004 | 0.003 | 0.194 |
| *Hamiltonella defensa* | -0.002 | -0.004 | 0.003 | 0.521 |
| *Regiella insecticola* | -0.000 | -0.004 | 0.003 | 0.225 |
| *Fukatsuia symbiotica* | -0.000 | -0.004 | 0.003 | 0.141 |
| *Ricketsiella spp.* | -0.002 | -0.004 | 0.003 | 0.445 |
| Co-infection: *H. defensa* + *R. insecticola* | -0.002 | -0.004 | 0.003 | 0.447 |
| **Beetle microbiome: Bottom-up models (both seasons)** | | | | |
| Microbiome diversity | -0.003 | -0.014 | 0.010 | 0.289 |
| Microbiome dominance | -0.008 | -0.141 | 0.010 | 0.606 |
